# Thermodynamically Explicit Kinetics with Potential for Genome-wide Application

**DOI:** 10.1101/2025.03.02.641056

**Authors:** Oliver Bodeit, Chalermroj Sutthaphirom, Emirhan Cenk Ulas, Rune Linding, Pimchai Chaiyen, Edda Klipp

## Abstract

Metabolic processes are inherently dynamic, yet genome-scale models are often limited to steady-state analysis, which cannot capture time-dependent responses crucial for understanding complex diseases like cancer. This work introduces a Thermodynamically Explicit (TDE) kinetic framework to bridge this gap by constructing dynamic metabolic models grounded in fundamental thermodynamic principles.

Our approach uses standard chemical potentials to derive simplified rate laws for metabolic reactions, reducing the kinetic complexity of each reaction to a single, biochemically determined prefactor (*λ*), which we will name kinetic capacity factor. The number of parameters in the suggested TDE kinetics is indeed minimal in the sense that it uses the lowest possible number of free parameters required to define a reaction rate that is both dynamic and thermodynamically consistent. We demonstrate the validity and practical application of this framework by re-parameterizing a well-established kinetic model of glycolysis. The resulting TDE model successfully reproduces the dynamic behavior of the original, more complex model in simulations.

By streamlining the parameterization process, the TDE kinetic framework offers a powerful and scalable tool for building genome-wide dynamic metabolic models. This approach paves the way for more accurate and predictive simulations of metabolic behavior, with significant potential for applications in systems biology, medicine, and biotechnology.

## 1 Introduction

### 1.1 Mathematical Modeling of Metabolism: From Steady-State to Dynamic Approaches

Metabolism is crucial to all life, and understanding its complexities is vital for various biological applications. Here, we focus on metabolic dynamics aiming to bridge the gap between steady-state and dynamic models. While genome-scale metabolic network reconstructions and constraint-based modeling like Flux Balance Analysis (FBA) have advanced our understanding of steady-state metabolic fluxes, their inherent limitation in capturing the dynamic nature of metabolic processes hinders our ability to predict and control metabolic behavior under changing conditions, especially in disease states like cancer.

Life depends on energy exchange, governed by the fundamental laws of thermodynamics. These laws are essential for comprehending the potential and limitations of cellular growth and metabolism. Systems biology utilizes mathematical models to describe cellular processes, but their predictive power relies on adhering to these fundamental physical and chemical laws, particularly thermodynamics. Life is inherently dynamic across all scales. Mathematical models explain observed behaviors in diverse biological processes, including metabolism. Notably, many metabolic phenomena can be explained by steady-state models, forming the basis of FBA and related approaches. However, these methods neglect temporal dynamics and quantitative values of metabolite concentrations, both crucial for deeper understanding of biological processes. Non-equilibrium thermodynamics provides the framework to address time-dependent processes, determine directionality, and quantify energy expenditure associated with dynamic trajectories.

### 1.2 Introducing Thermodynamically Explicit (TDE) Kinetics: A Novel Approach

To bridge the gap between steady-state and dynamic models, we propose integrating thermodynamic principles into kinetic models for enzymatic reactions. We present a Thermodynamically Explicit (TDE) framework to transform existing FBA models into dynamic and quantitative models of metabolism. This approach aims to be mathematically robust, computationally tractable, and widely applicable. TDE kinetics offers mathematical correctness, simplicity in obtaining or estimating parameters compared to classical kinetics, and utility in parametrization of large metabolic networks.

In this work, we will first revisit fundamental principles starting with phenomenological equilibrium thermodynamics, progressing to open systems, steady states, and deviations. On this basis, we will, second, introduce the concept of TDE kinetics and the relevant mathematical formulations. Third, we will apply the approach to an established, carefully parameterized model of glycolysis in order to demonstrate the practical application and also the challenges coming with the new approach. Eventually, this research aims to develop a novel, thermodynamically grounded approach to model metabolic dynamics, enhancing our understanding and predictive capabilities beyond the limitations of current steady-state methods. We foresee that applications will include biotechnological processes, such as chemostat and batch cultures, as well as the metabolic shifts observed between healthy and diseased cells. A conceptual overview is provided in Figure 1.

## 2 The Thermodynamically Explicit (TDE) Kinetics Framework

### 2.1 From Mass Action To Thermodynamics

For the classical description of the dynamics of metabolic networks, a number of kinetic rate laws have been introduced. With a few exceptions, they are all based on the law of mass action introduced already in the 1860s by Guldberg and Waage [GW67, WG64]. This law states that a reaction rate *v* is proportional to the rate of collision of the reactants which, in turn, is proportional to their concentration to the power of the molecularity with which they enter the reaction. Exemplified for the following reaction

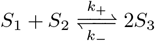

the reaction rate would read

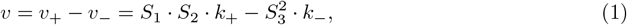

where *S*_1_, *S*_2_, and *S*_3_ also denote the concentrations of *S*_1_, *S*_2_, and *S*_3_, resp., and *k*_+_ and *k*_−_ are the forward and backward kinetic constants, respectively.

For enzyme catalyzed reactions, more complex mechanisms have been formulated, especially the well-known Michaelis-Menten rate equation, [MM^+^13]) with parameters *V*_*max*_ for each forward and backward reaction and Michaelis-Menten constant *K*_*M*_ for each metabolite in each reaction. Various rate equations have been introduced that take different mechanisms for saturation, inhibition, ligand binding or cooperativity into account. For reactions with two or more substrates there are, for example, ordered bi-bi mechanism, ping-pong kinetics, random kinetics. To simplify and generalize these various kinetics, we introduced the so-called convenience kinetics [LK06], which is now frequently used especially in computational approaches. Here, we come back to the law of mass action to consistently employ thermodynamics.

**Figure 1.**
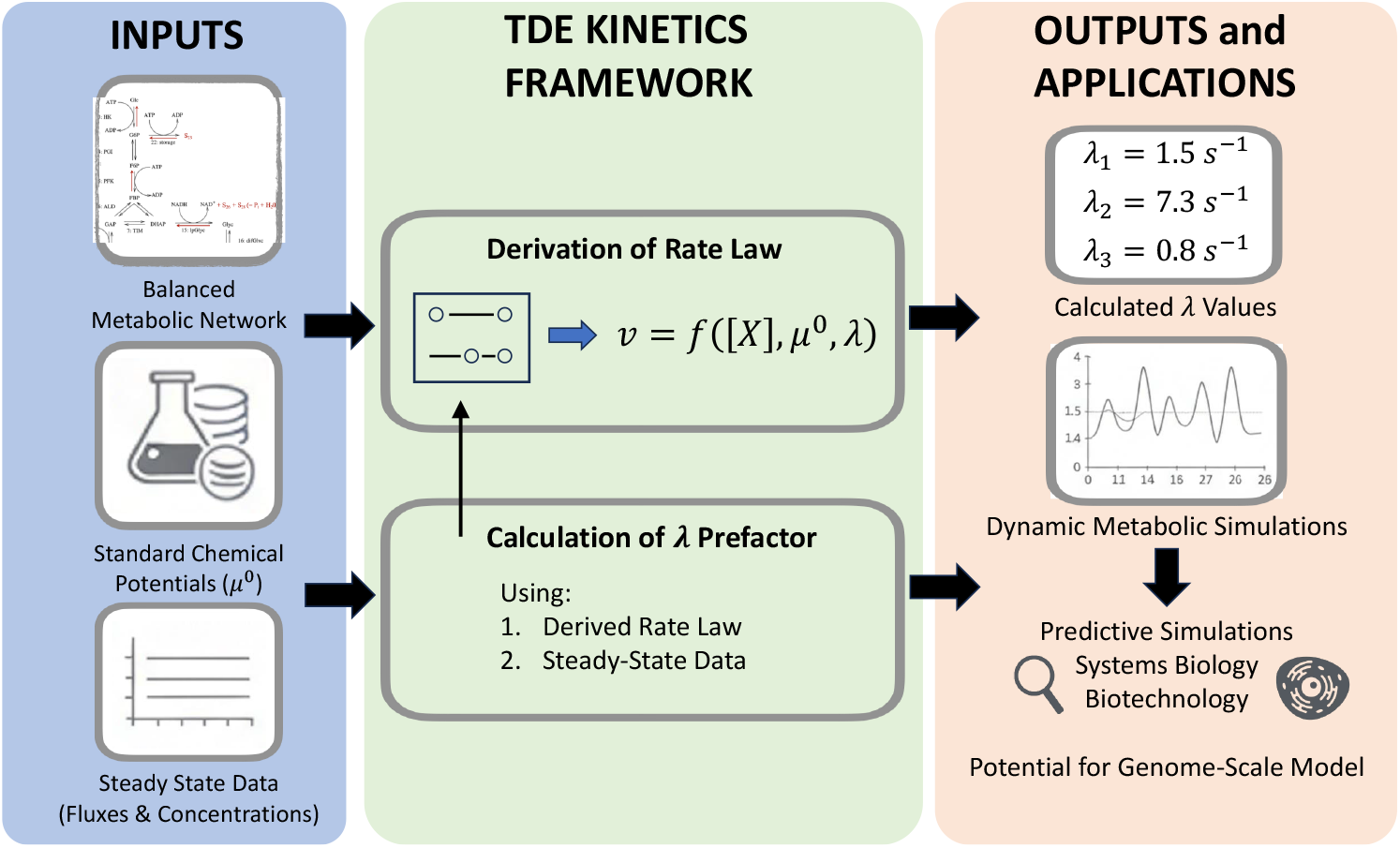
Overview over the concept of Thermodynamically Explicit Kinetics. The required inputs are (i) a precisely balanced metabolic network, i.e. all reactants must be listet, (ii) the standard chemical potentials of the reactants, and (iii) a list of steady state data for reaction fluxes and reactant concentrations, that is preferentially but necessarily complete (limited lack of data can be dealt with in algorithmic approaches). The TDE Kinetics framework provides the means to calculate the only missing parameter per reaction, i.e. the *λ*-value, based on the provided data. For the case of missing data, an optimization algorithm can be used to estimate those. The result is a parameterized kinetic model without thermodynamic inconsistencies for the metabolic network, that can be used for dynamic simulations and quantitative analysis.

### 2.2 Derivation of the TDE Rate Law

To motivate the TDE rate law and outline its relation to classical kinetics, let’s start with considering a single monomolecular reaction *r* with one substrate, *S*_*i*_, and one product, *S*_*j*_.

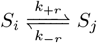

where *k*_+*r*_ and *k*_−*r*_ are the forward and backward kinetic constants, respectively. Be it further assumed that all reaction rates *v* follow mass action kinetics (i.e. *v*_+*r*_ = *S*_*i*_ · *k*_+*r*_). Then, the reaction rate reads

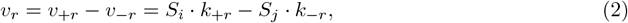

where *S*_*i*_ and *S*_*j*_ also denote the concentrations of *S*_*i*_ and *S*_*j*_, resp.. The chemical affinity *A*_*r*_ of the reaction is defined as

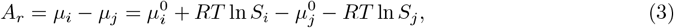

with *µ*_*i,j*_ being the chemical potential of compounds *S*_*i,j*_ and 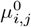 its standard chemical potential, respectively. *R* is the gas constant and *T* temperature. Note that under standard conditions (all concentrations measured in 1 M), the affinity *A*_*r*_ equals the negative value of the Gibbs free energy as also discussed below in the application. At equilibrium, both affinity and reaction rate vanish, i.e. it holds that *A*_*r*_ = 0 and, thus,

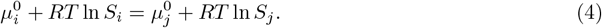

Rearranging of (4), while considering that also *v*_*r*_ = 0 at equilibrium, results in

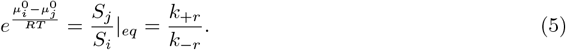

Equation (5) provides us with a relation between the rate constants *k*_*±r*_ and the chemical standard potentials *µ*^0^,

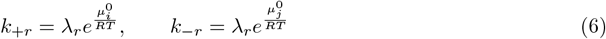

with a yet undefined parameter *λ*_*r*_. Since both the chemical standard potentials and the kinetic constants are constants, this relationship holds also away from the equilibrium (although we derived it for the equilibrium). The parameter *λ*_*r*_ results from the fact that for the reaction rate only the difference in chemical potentials of the reactants matters, not their absolute values. *λ*_*r*_ can be interpreted as being related to the height of the activation energy along the reaction coordinate and is dependent on both the amount and the activity of the catalyzing enzyme. Therefore, we name it kinetic capacity factor *λ*_*r*_. Note that the transition from Equation (5) to Equation (6) is not unambiguous in the mathematical sense, meaning that with Equation (6) also 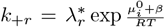 and 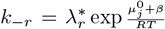 with 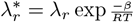 for every real number *β* (in appropriate units) would also be a valid solution. We will use this fact in the application below.

Introducing expression (6) into the general rate equation (2), the reaction rate *v*_*r*_ reads

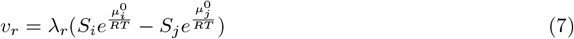

This expression derived here for the rate of an isolated reaction, will now be extended for biochemical reaction networks.

### 2.3 Network-Level Formulation of TDE Kinetics

In biochemical reaction networks with *R* reactions and *K* reactants, reactants can undergo different reactions at the same time. This is typically expressed by balance equations for their concentration changes either separately for the *i*-th compound as

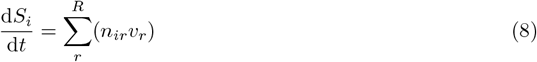

or in matrix form as

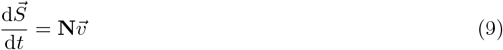

where

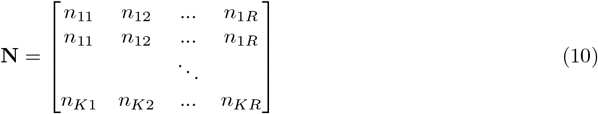

is the stoichiometric matrix of the reaction system.

For the purpose of relating kinetic parameters of individual reactions to the network stoichiometry, we will below separate the stoichiometric matrix given in Equation (10) into a producing part and a degrading part

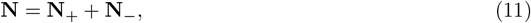

where **N**_+_ comprises only stoichiometric coefficients for the production of reactants (*n*_*ij*_ *>* 0) and **N**_−_ only for the degrading contributions (*n*_*ij*_ *<* 0). Below, they will together be called the partial stoichiometric matrices. We use

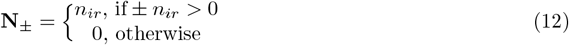

Let’s consider an example. For the following network consisting of three reactants and two monomolecular reactions

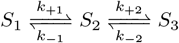

the full stoichiometric matrix as well as its degrading and producing parts read

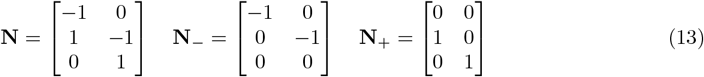

Next, we create two new vectors, 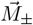, by multiplication of the vector containing the standard chemical potentials, i.e. 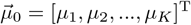 with the partial stoichiometric matrices, i.e.

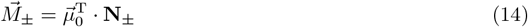

Here, the superscript T denotes the transpose of the vector (i.e. switch between a row and a column vector). Then, the kinetic constants of the *r*-th reaction read as follows

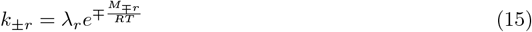

where *M*_*±r*_ is the *r*-th entry of 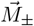.

For the above example (two successive monomolecular reactions), these quantities read

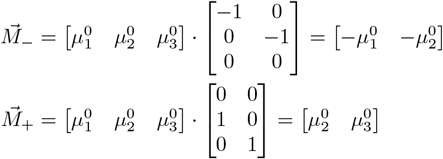

and

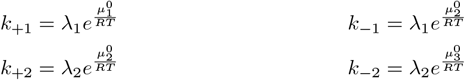

For a bimolecular reaction, exemplified by the following reaction scheme,

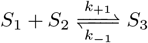

with the following set of stoichiometric matrices

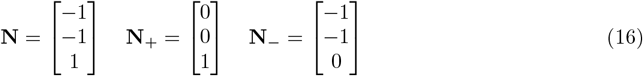

we would obtain the affinity

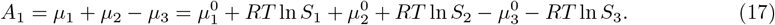

The equilibrium condition, with *S*_1_, *S*_2_ *≥* 0, then reads

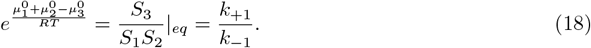

The 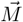 -vectors have only one entry each and read

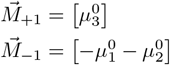

Consequently, the kinetic constants for this reactions are given by

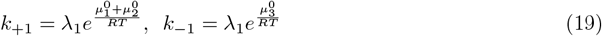

We note that all standard chemical potentials of the substrates enter a sum in the first exponential function. The same would hold for products and the second exponential, if there would be more than one product.

Thus, the general expression for kinetic constants of the *r*-th reaction read as follows

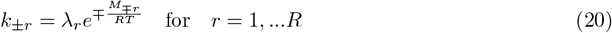

where *M*_*±r*_ is the *r*-th entry of 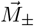.

Using the partial stoichiometric matrices **N**_+_ and **N**_−_ we calculate the rates for all reactions as follows

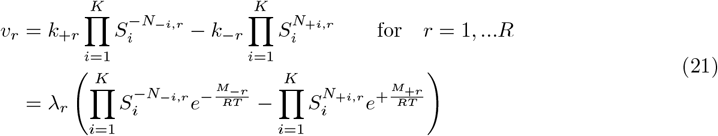

where *N*_*±i,r*_ denotes the (*i, r*)-th entry of the positive or negative partial stoichiometric matrix of **N**, respectively.

A matrix formulation for networks with only monomolecular reactions is provided in the appendix.

## 3 Methods: Parameterizing a Glycolysis Model with TDE Kinetics

### 3.1 Case Study: The Hynne Model

In order to demonstrate that the suggested kinetics is applicable to metabolic networks, we used an introduced and frequently used model of central carbon metabolism in yeast, the Hynne model [HDS01]. It was developed to explain oscillatory behavior after addition of cyanide. It contains 22 metabolites and 24 metabolic reactions, the kinetics and parameter values of which have been carefully extracted from literature and adapted to experimental data.

In applying the TDE kinetics to this model, now considered as ground truth, a series of practical problems have to be solved and technical considerations have to be taken. First, we need to obtain values for the standard chemical potentials for all represented metabolites and ensure that all model compounds are used in the appropriate physical units. Second, we have to revisit the stoichiometric matrix to ensure that all thermodynamically relevant metabolites are appropriately represented and that explicit and implicit model assumptions (e.g. constant values for external metabolites) are taken into account. Third, we have to estimate the values of kinetic capacity factors 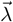. And eventually, we have to test in simulations that the resulting new model shows comparable dynamics to the original model.

To guide the reader’s imaging about the model used, we provide a scheme of the reaction network containing both the original version and our additions (as discussed below) in Fig. 2. More information about the model can also be retrieved from the JWS online database [PEVN^+^17] containing implemented and simulatable models of many metabolic and other networks under https://jjj.mib.ac.uk/models/hynne/simulate/.

### 3.2 Model Curation and Data Acquisition

#### 3.2.1 Obtaining Standard Chemical Potentials

The network-based parametrization of metabolic reactions requires two types of parameters: the standard chemical potentials 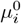 of each metabolite and the kinetic capacities or prefactors *λ*_*r*_ for each reaction. While classical chemical and biochemical literature provides numerous Gibbs energies for the reactions some standard chemical potentials (often calculated from formation enthalpies), a systematically collected and inherently coherent set of values is needed. Critically and as discussed above, all rates depend on *differences* of standard chemical potentials, not on absolute values.

To obtain a potentially genome-wide collection of standard chemical potentials, we used the publicly available tool eQuilibrator [BGM^+^22]. This tool employs a group contribution method [Alb94, Alb03, Mav91, NHMF13] to calculate reaction free energies, providing coherent values for user-selectable physical conditions like pH value and ionic strength. It also offers a comprehensive explanation of relevant thermodynamic terms (e.g. Gibbs free energy of reactions, Gibbs energy of formation, standard conditions), which we briefly summarize it here, aligning it with our notation.

“Standard conditions” imply all substrates and products are at 1 M concentration, denoted by a degree symbol “°” as in 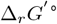 for the Gibbs free energy of the *r*-th reaction. Standard conditions serve as reference point, as reactions rarely take place under these conditions. Concentration changes within the reaction system affect 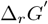, which represents our concentration dependent affinity *A*_*r*_. Realistic biological concentrations can shift 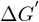 (or *A*_*r*_) from 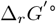 by tens of kJ/mol.

Gibbs free energy quantifies the work available within a system at constant temperature and pressure. eQuilibrator assumes that a temperature of 25°C (298.15 K) and a pressure of 1 bar. Δ_*r*_*G*° presents the change in Gibbs free energy for a reaction under standard conditions, excluding effects of pH, ionic strength or any other cellular factors (assumed to be zero). 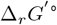 represents the change in Gibbs free energy at a specific pH and ionic strength. Neither Δ_*r*_*G*° nor 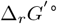 accounts for reactant concentration effects on free energy.

**Figure 2.**
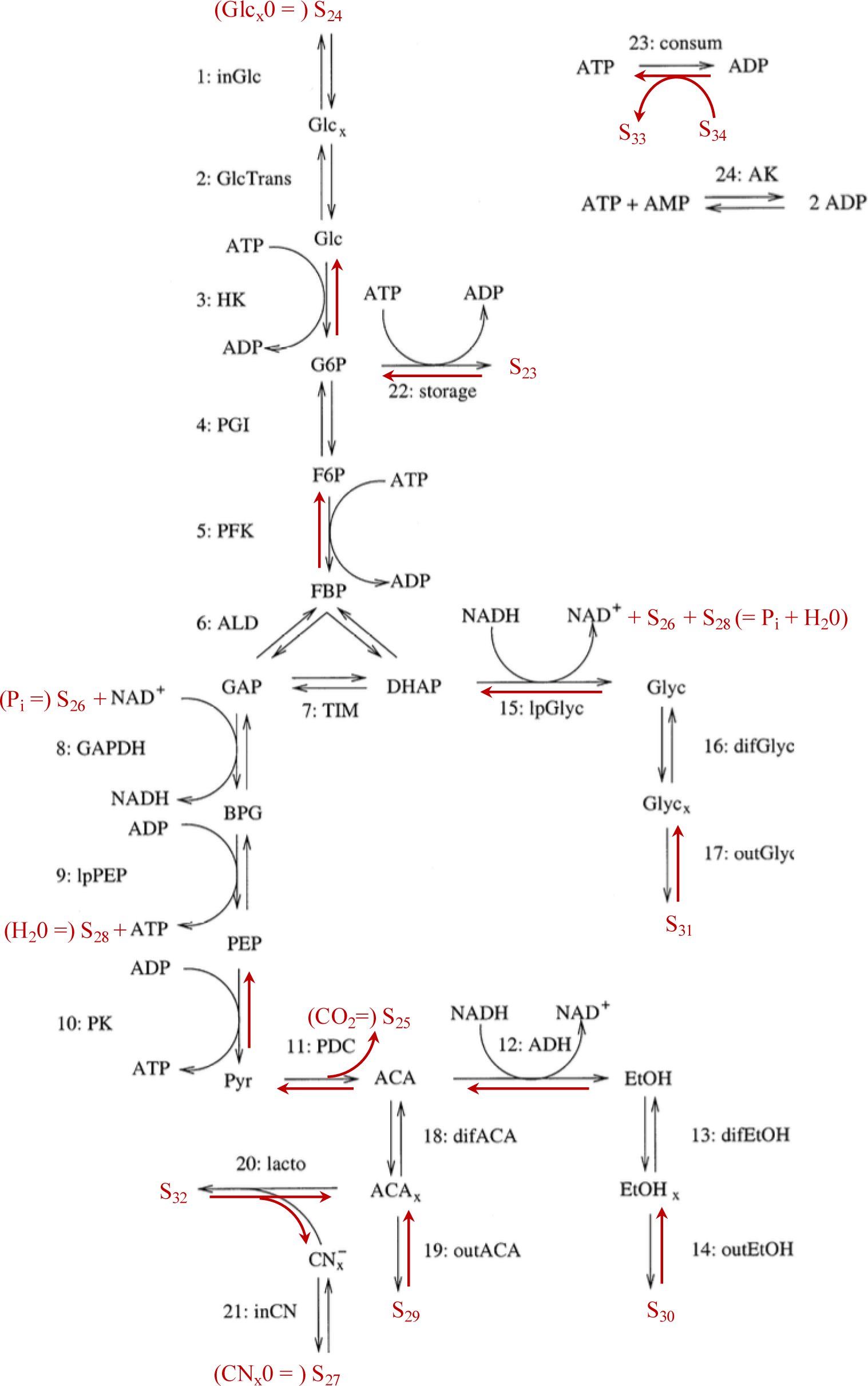
Adapted Hynne model. Reaction network of the Hynne model (adapted from [HDS01]). Black arrows, compounds and reaction annotations are original. Red arrows and compounds have been added here for correct reaction balancing.

To incorporate concentration effects, we decomposed the chemical affinity *A*_*r*_ into the standard chemical potential and a term dependent on the logarithm of the metabolite concentration (in M), as shown in Equation 3. The standard chemical potential reflects the energy state defined by a compound’s formation energy—the change in free energy during its formation from elements in their standard state. In eQuilibrator, Δ_*f*_ *G′* ° is the formation energy of a compound under specific cellular conditions defined by pH and ionic strength. The default pH is 7 and the default ionic strength is 0.1 M.

For details on how these different values are incorporated into the group contribution method, refer to eQuilibrator and to [NBEF^+^12]. eQuilibrator also provides free energy values adapted to 1 mM concentration (Δ_*r*_*G′m*, Δ_*f*_ *G′m*), recognizing that this is often more representative of metabolic concentration. While we used Molar units for all concentrations and parameters, using mM would be equally valid.

To parameterize the Hynne model with the TDE kinetics, we used Δ_*f*_ *G′* ° from eQuilibrator at pH 7.5, pMg 3.0, and 0.25 M ionic strength. All values are given in Table 2 for traceability. However, applying these values to a model with realistic concentrations and rates revealed two unit-related problems, discussed below.

#### 3.2.2 Defining Distinct Stoichiometric Matrices to Account for Sinks and Volume Differences

The choice to use Δ_*f*_ *G′* °, i.e. those values determined for standard conditions requires also to adjust all concentration and parameter values of the Hynne model respectively. That means that all concentrations have to be used in M (or practically multiply all unit-less given values in JWS online by 10^−3^). Equally, all Michaelis-Menten constants, maximal velocities and other rate constants have to be considered, if they contain the unit of concentration.

Another issue is that the value of gas constant multiplied with temperature of about *RT* = 2.47897 kJ/mol is small compared to many values of standard chemical potentials. Therefore, the practically resulting value of 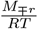 needed to calculate the parameter values (e.g. Equation 15) can be in the order of a thousand, potentially resulting in numerical issues, if we want to calculate 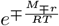. Since the assignment of all *µ*^0^s of substrates to *k*_+_ and all *µ*^0^s of products to *k*_−_ at the transition from Equation 5 to 6 is only determined up to a linear factor (the value *β*), we can inflict any factor to both kinetic constants without violation of thermodynamic constraints (though change of *λ*_*r*_). Different choices are possible as long as Equation 5 is fulfilled. We have chosen to subtract the mean of *M*_+*r*_ and *M*_−*r*_ (*β* = − (*M*_+*r*_ + *M*_−*r*_)*/*2) from both of them for each reaction. This results in values relatively close to zero, such that the calculation of the exponential function does not create numerical issues.

#### 3.2.3 Revising the Stoichiometric Matrix N to Ensure Thermodynamic Balancing by Adding of Omitted Metabolites

The publication of the Hynne model does not provide its stoichiometric matrix **N** directly, but it is evident from the provided set of reactions (their Table 1 and Figure 2). **N** could also be retrieved from the JWS online database. However, we cannot use this matrix directly for a number of reasons.

First, the Hynne model has omitted a number of metabolites that are critical for correct balancing of Gibbs free energies of reactions. Note that we have numbered metabolites included in the Hynne model according to their appearance in tables (see Table 2). Then, we have added the missing metabolites as follows:

- Reaction 11 (PDC) lacks CO_2_, added as *S*_25_
- Reaction 9 (lpPEP) lacks water, added as *S*_28_
- Reaction 8 (GAPDH) lacks orthophosphate, added as *S*_26_
- Reaction 15 (lpGlyc) lacks both water and orthophosphate, added as *S*_28_ and *S*_26_

Second, the implementation on JWS online has included the difference in volume between inside the cell and the external medium as a value of 59 in the respective uptake or release reactions, in accordance with chosen value by Hynne et al. We adopted this change, however only for the calculation of concentration changes, not for the calculation of *λ*_*r*_-values or reaction rates as detailed below.

Third, since our approach considers all reactions as basically reversible, we also added separate metabolites for the sinks of outgoing reactions (there called “P”), i.e.

- Reaction 14 (outEtOH) gets *S*_30_ as sink
- Reaction 17 (outGlyc) goes to *S*_31_
- Reaction 19 (outACA) is supplemented with *S*_29_

The concentrations of the added external compounds will still be considered as vanishing and fixed. Fourth, reactions 22 (“storage”) and 23 (“consum”) lump many processes consuming glucose-6-phosphate (22) and ATP (22 and 23). Hence, they are also not balanced, which would lead to unrealistic values of Gibbs free energy. In order to adjust the model respectively, we added another product to reaction 22 (*S*_23_) and substrate and product to reaction 23 (i.e. *S*_33_ and *S*_34_). In addition, the fixed external species (*Glc*_*x*_0 = *S*_24_ and *CN*_*x*_0 = *S*_27_) need to be taken into account.

These adjustments lead to two different types of stoichiometric matrices, called here **N**^*λ*^ and **N**^*v*^ (Equation 7.1 in Appendix). Both matrices contain 24 columns for the reactions considered in the Hynne model. Deviating from the Hynne model, they contain in total 34 rows for all metabolite listed above, i.e. the 22 original metabolites of the Hynne model, as well as the added metabolites to complete the reactions correctly, the sinks for excreted compounds, and the dummy metabolites for lumped processes. **N**^*λ*^ is used, first, to calculate the kinetic constants via its partial stoichiometric matrices and the 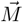 vector as exemplified in Equations 11 to 15. Here, it reads

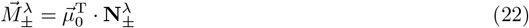

Then, the kinetic constants of the *r*-th reaction read as follows

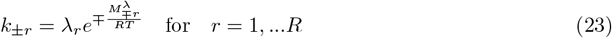

where 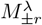 is the *r*-th entry of 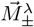.

Second, the matrix **N**^*λ*^ is used to calculate the rates for all reactions as follows

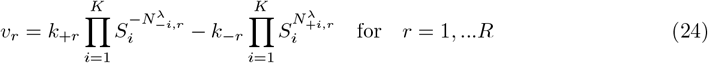

where 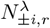 denotes the (*i, r*)-th entry of the positive or negative partial stoichiometric matrix of **N**^*λ*^, respectively.

The matrix **N**^*v*^ has two types of modification compared to the matrix **N**^*λ*^. First, all entries in all rows for added metabolites (here in rows 23 to 34) are set to zero to ensure that those metabolites are kept constant in dynamic simulations. Second, matrix **N**^*v*^ is modified at the entries that reflect transport between different volumes where the respective entries are set to +59 or -59 as in JWSonline. In a general case, these values should reflect the volume ratio between the two compartments that are source or sink for the transport reaction. They should not be confused with stoichiometric coefficients in the sense of numbers of metabolites needing to collide for the reaction to happen. **N**^*v*^ is used calculate the concentration changes during temporal simulations as in Equation 9.

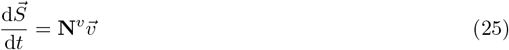

Equations 22 to 25 are now the full set of equations required to simulate the dynamic model given the values for the standard chemical potentials and initial concentrations, however, with still unknown parameters *λ*_*r*_. Next, we discuss how to obtain *λ*_*r*_-values.

### 3.3 Calculation of the Kinetic Capacity Factors *λ*_*r*_

If one intends to formulate a dynamic model for a metabolic network without prior knowledge, *λ*-values need to be estimated from data for metabolite concentrations and fluxes, as in classical parameter estimation problems. Here, we had the advantage of a fully parameterized model to be used as “ground truth” and, therefore, we used steady state fluxes and concentrations of the Hynne model to derive *λ*-values for all reactions and then confirmed their suitability by simulating and comparing the dynamic behavior starting with different initial conditions (metabolite concentrations) using either the published or TDE kinetics. Since *λ*-values should be independent of the specific state (i.e. steady state or not), we rearrange Equations 23 and 24 to yield the kinetic capacity factor

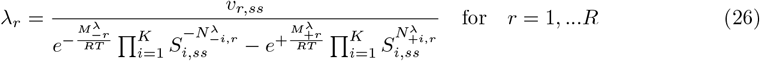

where *v*_*r,ss*_ and *S*_*i,ss*_ are reaction rates and metabolite concentrations in steady state, respectively. The resulting values for *λ*_*r*_-values for all reactions are given in Table 3 in the Appendix. Below, we will use these values to simulate the Hynne model and assess the sensitivity.

## 4 Results

### 4.1 TDE Kinetics Accurately Replicates Classical Dynamics

#### 4.1.1 Correlation of Individual Rates

To analyze the behavior of the kinetics under different conditions, we need a systematic comparison of the TDE kinetics with classical kinetics. Here, we look at all the reactions that are present in the Hynne model. They comprise simple irreversible mass action (e.g. reactions 14, 17, 19, 20, 22, 23), reversible mass action (reactions 1, 9, 13, 16, 18, 21, 24), irreversible and reversible Michaelis-Menten kinetics with different degrees of inhibition (reactions 2-4, 6-8, 10-12, 15) as well as an irreversible Hill kinetic (reaction 5). The TDE kinetics instead is in all cases a reversible mass action kinetics. Therefore, it is not self-evident that the behavior should reflect that of classical kinetics. As shown in Fig. 3, we have tested that as follows: we have chosen a steady state with external glucose concentration *Glc*_*x*_0 = 5 · 10^−3^M and external cyanide concentration 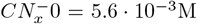 as reference point. For this point, we calculated *λ*_*r*_ as explained above. Then, we sampled the concentration space around the steady state concentration one thousand times choosing each time a new set of random values for the concentrations between their 0.5- and 1.5-fold (uniform distribution). For these sets, we calculated all the rates either with the classical kinetics used by Hynne et al. or with the TDE kinetics. As can be seen in Fig. 3, most rates show comparable values for both approaches; the correlation coefficients are given on top of the panel for each reaction. We see that all reactions that have mass action kinetics at Hynne display a correlation coefficient of 1, thus a replacement has no influence. An exception is reaction 24 which is forced to have zero rate in steady state (because AMP does not enter any other reaction) and therefore leads to numerical uncertainties in determining *λ*_24_. Many of the reactions with Michaelis-Menten kinetics at Hynne show correlation coefficients in the order of 0.9 and even above. Given that parameter estimation with real metabolites often very tedious and good fits are hard to achieve, this argues for TDE kinetics as an acceptable replacement. The only clear exception is reaction 5 with a correlation coefficient of only about 0.1. This indicates that the sigmoidal behavior provided by Hill kinetics is not optimally reflected with TDE kinetics, at least not on the level of the individual reaction.

#### 4.1.2 Sensitivity Towards Boundary Conditions

The *λ*_*r*_-values that have been calculated based on the steady state assumptions are not completely independent of the chosen boundary conditions affecting the steady state to be reached. In the Hynne model we can, for example, vary the concentration of external glucose (Glc_*x*_0), which leads to different steady states. Hence, also some of the *λ*_*r*_-values vary slightly as shown in in Fig. 4, however only to minor extent.

#### 4.1.3 Full Dynamic Simulation

With the preparatory steps discussed above, we are ready to simulate the Hynne model. As mentioned above, we simulated both the classical model and our newly parameterized TDE model for a set of example conditions to compare them. As reference point, we have chosen a steady state with external glucose concentration *Glc*_*x*_0 = 5 10^−3^M and external cyanide concentration 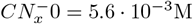. If we start directly at steady state, in both models the system remains in steady state, as required. If we perturb the initial concentrations, here by setting internal glucose to a hundredths of its original value (i.e. from *Glc*_*x*_ = 0.0801634 · 10^−3^M to *Glc*_*x*_ = 0.0801634 · 10^−5^M, the following dynamic behavior of the TDE model resembles the behavior of the classical model closely as seen in Fig. 5. There are, of course, small deviations, but the qualitative behavior remains.

**Figure 3.**
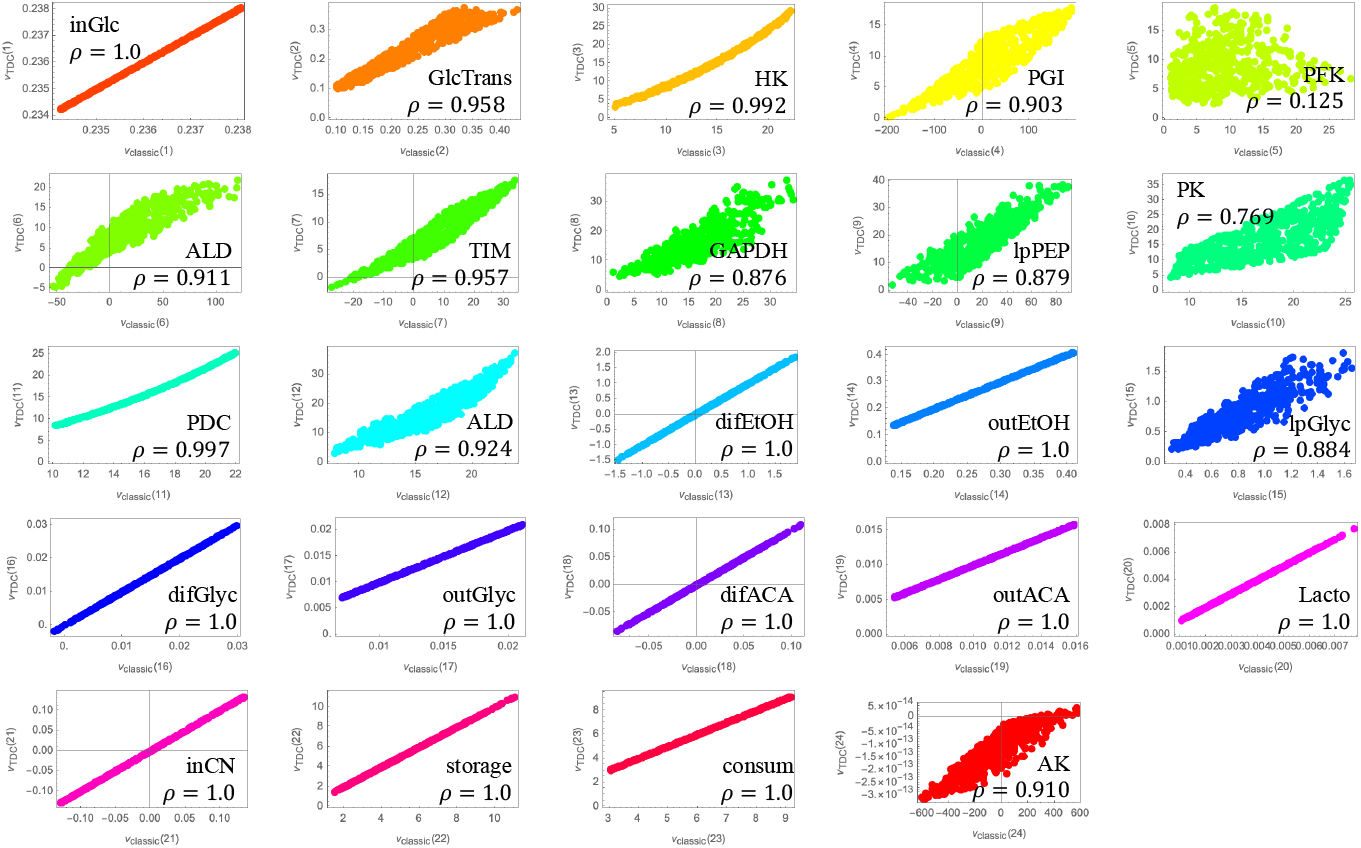
Comparison of classical kinetics to TDE kinetics for individual reactions. We created 1000 sets of metabolite concentrations varied randomly between the 0.5- and 1.5-fold of their steady state value and calculated both the classical rates according to the Hynne model and the TDE rates. Within the panel for each reaction is the correlation coefficient *ρ* is displayed, which is 1 for reactions with mass action kinetics in the Hynne model and around 0.9 for reactions with Michaelis-Menten kinetics. Only reaction 5 (PFK) with Hill kinetics shows deviating behavior.

**Figure 4.**
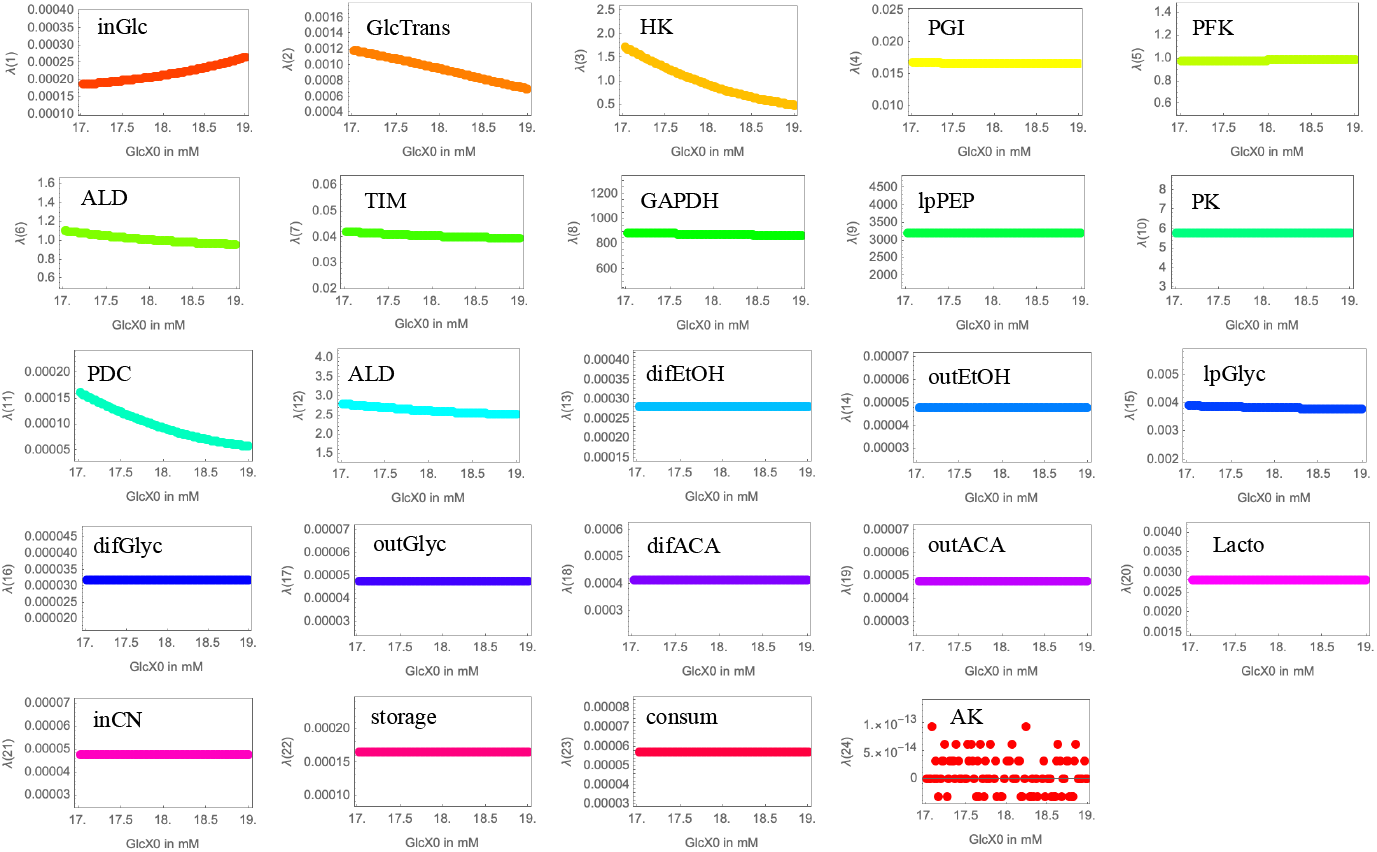
Dependence of kinetic capacities *λ* on boundary conditions. To test the effect of boundary conditions, we varied the external glucose concentration *Glc*_*x*_0 in a range between 17 · 10^−3^M and 19 · 10^−3^M, recalculated the steady state of the system and the resulting *λ*_*r*_-values. While *λ*_*r*_-values are constant for most reactions, few reactions show minor dependency.

**Figure 5.**
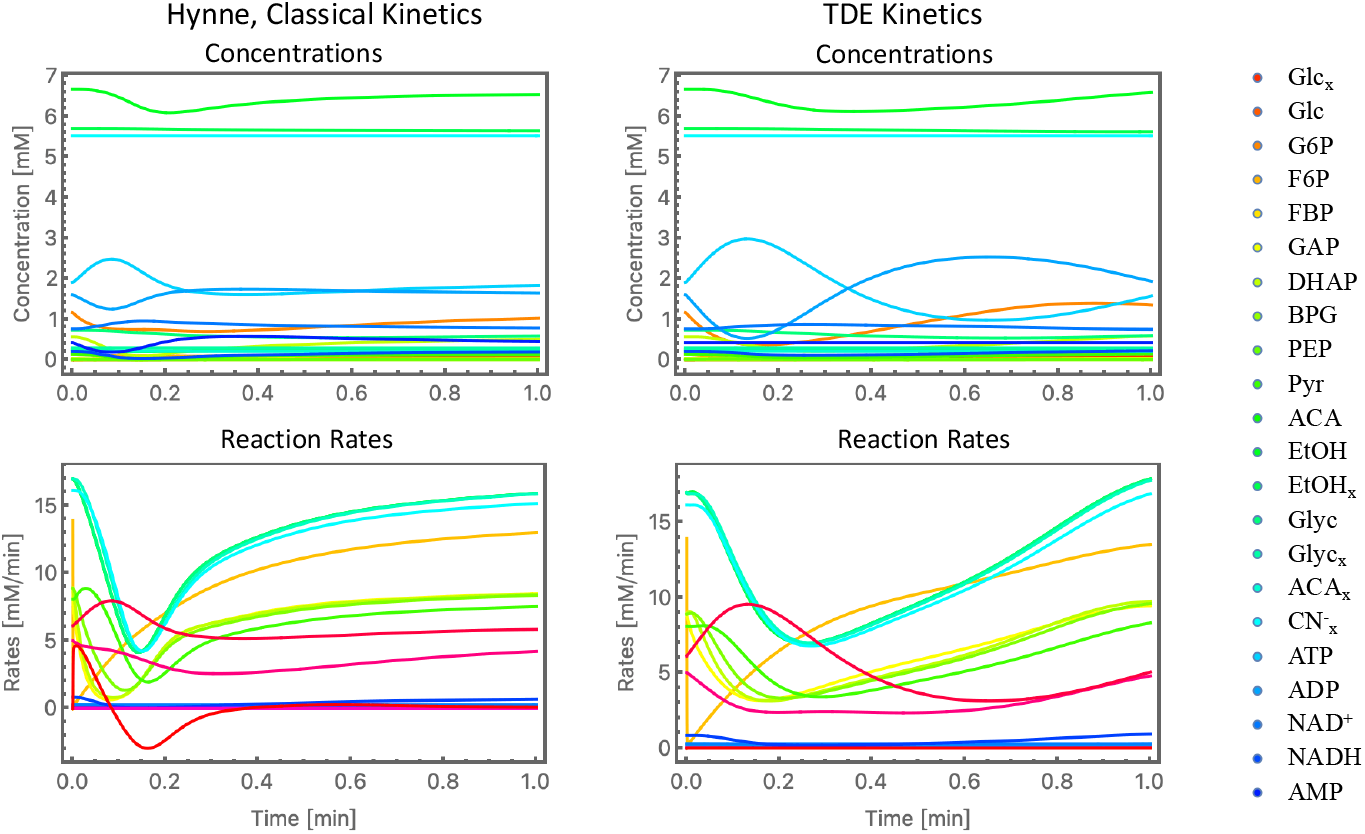
Comparison of model dynamics simulated with either classical kinetics or TDE kinetics. We perturbed the initial concentration of external glucose by setting it to its 0.01-fold compared to the steady state concentration. Left panels: Hynne original kinetics, right panels: TDE kinetics. Top panels: concentration time course, bottom panels: time courses of reaction rates. Right edge: color code for metabolites. Color codes for reactions see Fig. 6.

### 4.2 The Kinetic Capacity Factor *λ* Reflects Thermodynamic Constraints

#### 4.2.1 Correlation Between the Logarithm of *λ* and Sums of Standard Chemical Potentials of the Reactants

Having a deeper look into the properties of *λ*_*r*_-values, we find that the above calculated values show a tendency to become the larger the more negative the sum of the standard chemical potentials of all substrates and products of a reaction are. To that extend, we have plotted the natural logarithm of *λ*_*r*_-values against the sum of all standard chemical potentials of substrates and products of the *r*-the reaction, denoted here with *M*_+*r*_ + *M*_−*r*_, see Equations 14 and 15 and Figure 6. A linear fit of the type

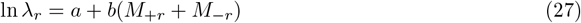

resulted in ln[*λ*_*r*_] = −1.4946 − 0.0019426(*M*_+*r*_ +*M*_−*r*_). We did not identify a similar relationship to the difference of *M*_+*r*_ and *M*_−*r*_, i.e. the free energy of the reaction. Figuratively speaking, the higher the activation energy of the reaction, i.e. the magnitude of the potential barrier separating minima of the potential energy surface pertaining to the initial and final thermodynamic state, the smaller the classical kinetic constants associated, but consequentially also the higher also the respective required kinetic capacity, i.e. the *λ*_*r*_-values, to enable the reaction flux. This may indicate that *λ*_*r*_-values are to some extend dictated by thermodynamic necessities and the cell has to provide catalysts that are respectively abundant and strong in their catalytic efficiency.

**Figure 6.**
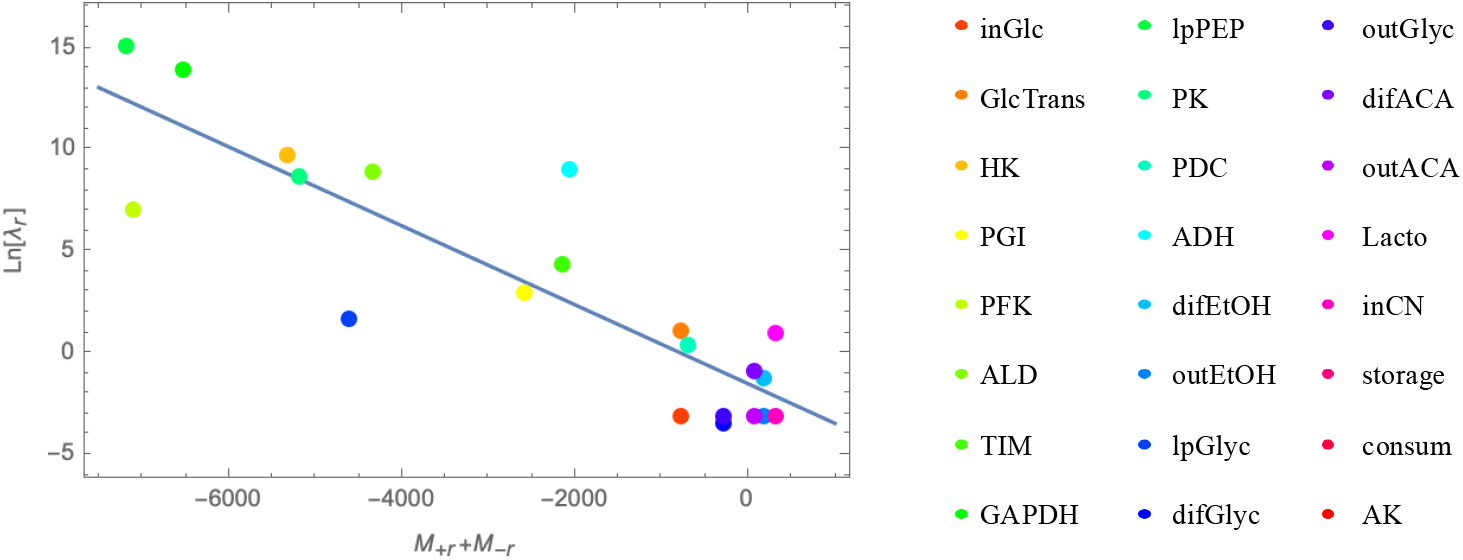
Relation between *λ* and *µ*^0^ values. The natural logarithm of *λ*-values is plotted against the sum of all standard chemical potentials of substrates and products of the respective reaction, denoted here with *M*_+*r*_ + *M*_−*r*_, see Equations 14 and 15. Reactions 22 to 24 have not been included, due to our arbitrary choice of parameters for added substrates. The gray line is a linear fit of data points to guide the eye. Right panel: color code for reactions valid for all figures.

#### 4.2.2 Correlation between the Kinetic Capacity Factor *λ* and *M* -Values in Other Models

We wanted to understand, whether this relationship between the natural logarithm of *λ*-values and the *M*_+_ +*M*_−_-values per reaction is just a feature of that specific Hynne model (and its underlying reaction system) or whether it reflects a general property of biochemical networks. Therefore, we analyzed a series of metabolic models, which are available in JWSonline and had been carefully curated before. The list of models together with our analysis results is given in Table 1.

**Table 1:**
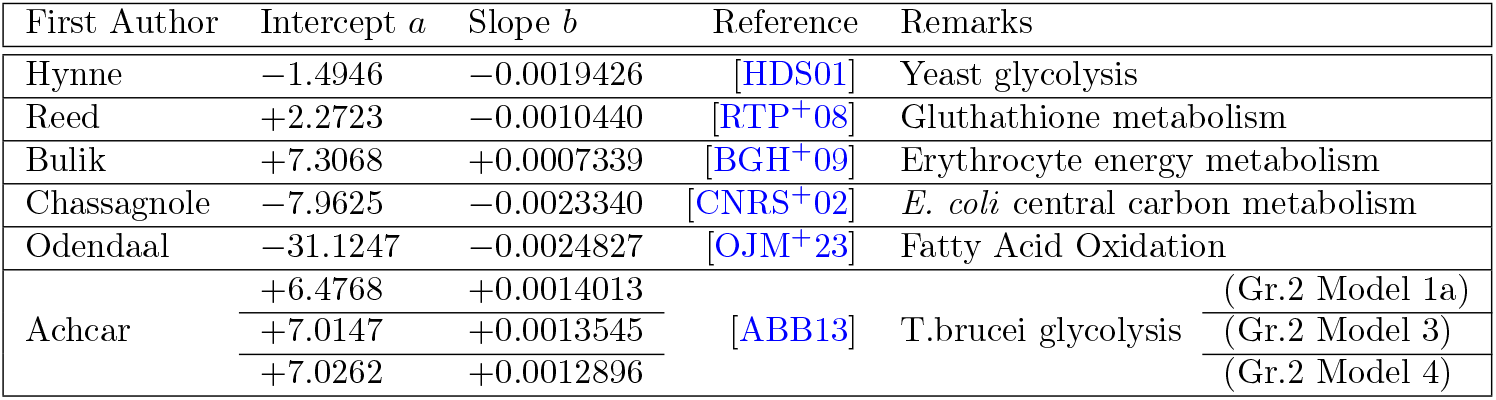
Models Analyzed with TDE Kinetics. Be the relation between ln *λ*_*r*_ and (*M*_+*r*_ + *M*_−*r*_) given by ln *λ*_*r*_ = *a* + *b*(*M*_+*r*_ + *M*_−*r*_) with *a* being the intercept and *b* the slope of the linear function. For more information see text.

All models had to be extensively revised by adding missing metabolites to ensure thermodynamic balancing. These were either internal metabolites (such as protons or water but sometimes also other omitted metabolites) or the external metabolites (which are often neglected or just considered as constant pool, but are relevant for our approach). Then, standard chemical potentials have been retrieved and *λ*-values have been calculated for steady states as described for the Hynne model above. Table 1 shows that the slopes are in about the same order of magnitude while the intercepts vary. Hence, the analysis of different models indicates that the specific relation can also be found in those models, but it doesn’t explain why.

#### 4.2.3 Application of Principles of Optimal Enzyme Distribution in Metabolic Networks

Let’s consult theory to further elucidate the finding. We had shown previously that biochemical reaction networks in optimal states exhibit a special relationship between enzyme concentrations and flux control coefficients [KH99]. Optimal states here mean that a given set of steady state fluxes is realized by a minimal amount of total enzyme concentration (assuming that enzyme concentrations could be freely assigned to different reactions or that evolution has lead to a state where the cell has to produce the least overall amount of enzymes that can realize a given flux pattern necessary for successful growth). Flux control coefficients represent a kind of normalized sensitivity coefficients of the steady state fluxes *J*_*j*_ with respect to the rates *v*_*r*_ of individual reactions. They are defined as

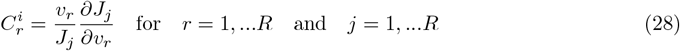

One can easily argue that in Eq. 28 the rate *v*_*r*_ can be replaced by the enzyme concentration *E*_*r*_ (as done in [KH99]) or by our newly introduced quantity *λ*_*r*_, because both *v*_*r*_ and *λ*_*r*_ are proportional to *E*_*r*_. The optimality condition derived in [KH99] reads

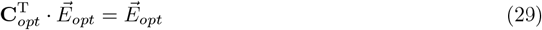

where **C**_*opt*_ is the *R × R* matrix of all flux control coefficients (all steady state fluxes versus all rates) in optimal states, T denotes the transpose, and 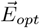 is the vector containing all optimal enzyme concentrations. Following [KH99] this expression is equivalent to the following expression

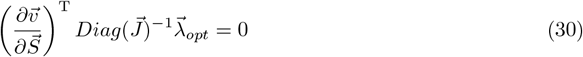

where we already replaced optimal enzyme concentrations by optimal *λ*-values. The matrix 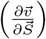 denotes the so-called elasticities, i.e. the partial derivatives of reaction rates w.r.t. metabolite concentrations, here to be evaluated at the optimal state. 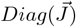 is the *R × R* matrix with steady fluxes in its main diagonal, resulting from the normalization used in Eq. 28. The expression 30 already establishes a relationship between *λ*-values and the kinetics of the network. Now, we need to evaluate how that relates to the dependency the logarithm of *λ*-values and the *M*-values.

To keep it simple, we will illustrate the consequences for *λ*-values only for a linear pathway as shown below

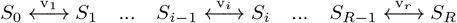

where *S*_0_ to *S*_*R*_ are the series of metabolites of the pathway. Here and below, all equations hold for all *i* = 1, …*R*. In the Appendix, we describe a pathway that can also have additional substrates and products for each reaction as a step of generalization.

For the linear pathway with only single substrates and products per reaction, we can then easily show that the ratio of two successive *λ*-values in the pathway reads

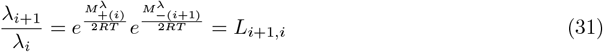

Here, we have already considered that the steady state fluxes are the same in all reactions of the linear chain and, hence, cancel out. Together with the condition of a fixed (minimal) total amount of *λ*-values, i.e.

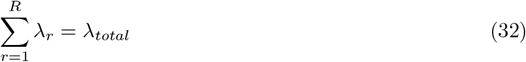

we easily find that

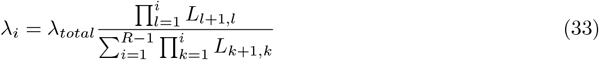

As an example, for a reaction chain with only three reactions and no further metabolites than the series of main metabolites (*S*_0_ to *S*_3_) such as

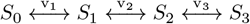

the natural logarithms of *λ*-values then read

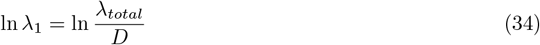

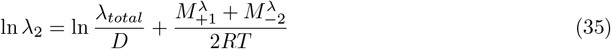

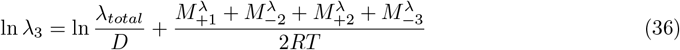

with

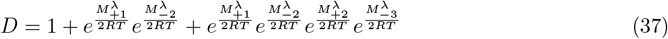

Let’s interpret this result. First, note that 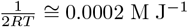 for room temperature (*T* = 300*K*), which is in the same order of magnitude as the slopes *b* listed in Table 1. Second, the factor behind *b* is, however, not directly equal to 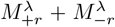, but instead a sum of *M*-values of the preceding reactions. This shows that, in optimal states, the required enzymatic activity represented here by *λ* is not only dependent on the actual reaction, but also on the activation energies of the upstream reactions. This resembles the fact that in linear pathways flux control is typically higher for reactions in the beginning than in the rear of the chain and that in optimal states enzyme concentrations should decline from the beginning to the end of the pathway (at least for the simplifying assumption that the kinetic parameters would be about the same in all reactions). In branched pathways or networks, the situation is, of course, more complicated and depends on both the network structure and the chemical potentials of reactants. Third, the intercept *a* can be clearly identified and reads here ln 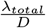. This means that *λ*_*total*_ dictates the order of magnitude of *λ*-values within the whole network. Like the total enzyme concentration discussed in the optimization approach, *λ*_*total*_ represents the metabolic capacity of the cell or its capacity to produce enzymes. One has to expect that this total capacity varies with external conditions, especially the availability of nutrients, but also potentially other factors such as cell type, cell function, or being either single (microbial) cell or part of a complex organism. Thus, for now *λ*_*total*_ is a quantity that has to be determined for each metabolic model with given organism and experimental conditions. *M*-values (and hence also *D*), however, result from the standard chemical potentials of the reactants and should be applicable to metabolic models for various organisms and conditions.

In the following, we will use this information to suggest an algorithm that supports the estimation of *λ*-values for models with only partially known rates and concentrations.

## 5 Estimation of *λ*-values for models with only partially known rates and concentrations

We developed a computational strategy that is an automated, iterative algorithm designed to calculate values of the kinetic capacities *λ* for metabolic systems with incomplete data. It requires knowledge of the reaction rates, thus sufficient information to perform flux balance analysis is essential. Then, in brief, we first use the available data to create a initial fit of type 27, which is, second, used to optimize the remaining *λ* parameters and missing metabolite concentrations. The process initiates by verifying that there is sufficient data before performing an initial “fit” calculation, which is afterwards refined by detecting and removing any data outliers. The quality of this initial fit is measured by its *R*^2^ value and dictates the subsequent steps. A high-quality fit (parameter 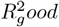 adjustable by the user, we used *R*^2^ *>* 0.5) is presented to the user for validation, while mediocre or poor fits trigger a more comprehensive optimization phase. This core optimization can be executed with “Standard Settings” or, if that fails to yield a good result, with “Higher Penalties”. A key feature of this stage is the application of a “Backtracking” procedure to refine the solution, with the algorithm checking if this step led to an improvement. When multiple potential solutions have been generated from different runs, the algorithm employs a global scoring system to select the best one. Points are awarded based on the *R*^2^ value, the slope of the fit, and whether the solution has many identical lambda values, which is penalized. The optimization run with the highest score is chosen as the final result, using tie-breaking rules if necessary.

In the Appendix, we provide an extended version of the algorithm and the resulting model improvements.

## 6 Discussion

We have introduced a thermodynamically explicit (TDE) kinetic approach that utilizes tabulated standard chemical potentials to derive individual kinetic parameters. The method separates the kinetics into a known, chemically determined component (the exponential function containing *µ*^0^) and a biochemically determined component *λ*-value, which also depends on enzyme amounts and properties. The resulting kinetics is essentially a mass-action kinetics. Comparable approaches have been discussed in the framework of bond graphs (e.g. in [GP22] with an application to photosynthesis) or in [Pfe24, EG07] studying toy models to explore the flux solution space. We have demonstrated how the stoichiometric matrix of the metabolic network under study can be used to systematically derive rate equations for the entire network. A key advantage of TDE parametrization is its potential application to metabolic networks of all sizes, from small to genome-scale. Given that eQuilibrator now contains data for approximately 100.000 compounds, providing impressive metabolic coverage, this extensive availability of *µ*^0^-values should facilitate TDE parametrization, particularly for large (potentially genome-wide) metabolic networks. However, it should be noted that for some complex molecules (like glycogen) or for artificially created molecules (i.e. binding of drug and metabolite) *µ*^0^-values are not immediately available.

Basic features of the TDE kinetics have been demonstrated using an accepted and well-studied model of yeast glycolysis introduced by Hynne et al. [HDS01], which employed different types of rate equation from mass action to Michaelis-Menten and Hill equation. To this end, we replaced all the classical kinetics with TDE kinetics, discussed the associated challenges and compared the resulting dynamics and steady state behavior with the original models. The comparison demonstrated that TDE kinetics is not a perfect copy of all classical kinetics, but can provide an easier way to the parametrization of the model. Our work with the Hynne model yielded several important insights. First, accurate balancing of all reaction substrates and products is crucial. Missing metabolites introduce errors in the sum of *µ*^0^-values and can lead to severe numerical issues. Fortunately, current community efforts for quality assurance in genome-scale modeling are strongly supportive to this requirement [LBO^+^20, JFC^+^25, TP10, SLZ^+^21]. Second, because only the difference of the sum of *µ*^0^-values of substrates and products are relevant, we subtracted their mean from both sums to avoid numerical issues. An alternative approach would be to set one sum to zero and the other to the difference. Third, we treated all reactions as, in principle, reversible. In practice, none of the reactions typically considered irreversible exhibited negative rates, also even in simulations not shown. Fourth, the original Hynne model had been parameterized to exhibit oscillations, a subject of in-depth analysis in their publication. Typical requirements for oscillations in a dynamic system include long-ranging feedback (which we preserved) and non-linear kinetics (which we largely removed). It is important to note that TDE kinetics is less effective at reproducing oscillatory behavior. Therefore, we selected example simulations within a range of boundary conditions that did not exhibit oscillations. Fifth, correct unit handling is essential. We chose to use Molar units for all concentrations and corresponding parameter values. An alternative, particularly using mM, would necessitate using the respective Δ_*f*_ *G′m*-values provided by, e.g., eQuilibrator.

Applying the TDE approach to a new model requires parameter estimation, specifically determining the kinetic capacities, i.e. the *λ*_*r*_-values. However, this involves only one parameter per reaction, unlike classical kinetics, which often require multiple parameters (e.g. *K*_*m*_, *V*_*max*_). A systematic approach would involve performing flux balance analysis to obtain steady state reaction rates, along with a set of steady state metabolite concentrations. Alternatively, time-resolved metabolite concentration data following a defined perturbation could also enable parameter estimation. It is worth noting that the reduction of unknown parameters to one parameter per reaction is the minimal set of parameters for that problem. That is also in line with previous deliberations of Ederer and Gilles, who used *R* + *K* parameters (*n* +*m* in their notation) [EG07]. If we would consider the standard chemical potentials as given, i.e. not as a parameter to be estimated, this number Would be reduced by *K* (or *m*). If we take into account the mutual dependency of steady state fluxes, we are left with a degree of freedom (and number of independently to be determined parameters) of *Rank***N**), i.e. the rank of the stoichiometric matrix.

For the general case of an unknown kinetic capacity *λ*, thermodynamically correct parametrization ensures that the resulting rate laws will not violate thermodynamic principles. However, it does not automatically guarantee that the parametrization accurately captures the reaction dynamics. Classical rate laws incorporate features such as dependence on enzyme activity (typically presented by enzyme concentration), saturation effects as well as activation or inhibition by other network components. These effects can be represented by an additional dimensionless prefactor for each affected reaction, as discussed for modular kinetic rate laws [LUK10].

In conclusion, we present a thermodynamically explicit parameterization of metabolic networks that requires a minimal set of parameters. This parameterization is, therefore, a candidate for deriving large-scale quantitative models of metabolism. An application to genome-scale models remains to performed since it requires a sufficiently large set of metabolomics data to perform flux analysis and the initial fit of the 27 relation.

## 7 Appendices

### 7.1 Stoichiometric matrices

The stoichiometric matrices for the revised Hynne model read

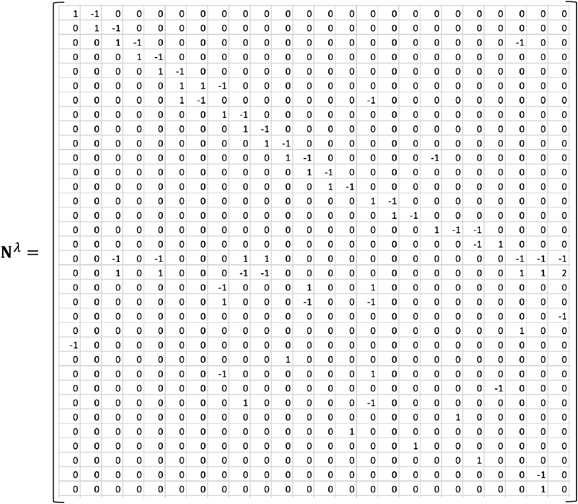

### 7.2 Metabolite and Reaction Information

Relevant numbering and quantitative information about metabolites and reactions in the Hynne model and our extension are provided in the tables below. Model results for the other models analyzed, i.e. stoichiometric matrices, given or optimized metabolite concentrations and fluxes as well as *λ*-values can be found in the Supplementary Table “Results of Model Analysis.xlsx”.

**Table 2:** Metabolite Information. Listed are all metabolites originally contained in the Hynne model (*S*_1_ to *S*_22_) with their steady state concentrations (for Glc_*x*_0 = 5.0 · 10^−3^) and their respective standard chemical potentials *µ*^0^ as well as those metabolites added for thermodynamic balancing (*S*_23_ to *S*_34_). Remarks: ^*^ *µ*^0^ was arbitrarily set to balance the differences in Gibbs free energy of the missing product. The concentration was set to zero (no re-influx). Note that to prevent numerical issues in Equations 24 and 26, i.e. to prevent 0^0^, a small non-zero value can be chosen instead. ^**^ concentrations were set to 1 to mirror their use in the Hynne model (multiplication by 1 does not affect the rate). ^***^ The concentration was set to zero (no re-influx). ^****^ concentrations were set to 1 to mirror their use in the Hynne model. *µ*^0^ was arbitrarily set to balance the differences in Gibbs free energy of the missing substrate and product. For more information see text.

| Number | Name | Concentration [M] | $\mu^0$ in [kJ/mol] | Remarks |
| --- | --- | --- | --- | --- |
| $S_1$ | $\text{Glc}_x$ | $8.01634 \cdot 10^{-5}$ | -392.3 | |
| $S_2$ | Glc | $7.29537 \cdot 10^{-6}$ | -392.3 | |
| $S_3$ | G6P | $1.16705 \cdot 10^{-3}$ | -1287.6 | |
| $S_4$ | F6P | $1.45314 \cdot 10^{-4}$ | -1285.0 | |
| $S_5$ | FBP | $1.92354 \cdot 10^{-4}$ | -2176.5 | |
| $S_6$ | GAP/G3P | $2.40510 \cdot 10^{-5}$ | -1074.4 | |
| $S_7$ | DHAP | $5.63423 \cdot 10^{-4}$ | -1080.1 | |
| $S_8$ | BGP/13DPG | $7.11072 \cdot 10^{-8}$ | -2194.9 | |
| $S_9$ | PEP | $1.15848 \cdot 10^{-5}$ | -1188.0 | |
| $S_{10}$ | Pyr | $1.41006 \cdot 10^{-4}$ | -338.1 | |
| $S_{11}$ | ACA | $2.54627 \cdot 10^{-4}$ | 34.3 | |
| $S_{12}$ | EtOH | $6.67191 \cdot 10^{-3}$ | 82.4 | |
| $S_{13}$ | $\text{EtOH}_x$ | $5.70552 \cdot 10^{-3}$ | 82.4 | |
| $S_{14}$ | Glyc | $7.30030 \cdot 10^{-4}$ | -145.0 | |
| $S_{15}$ | $\text{Glyc}_x$ | $2.93243 \cdot 10^{-4}$ | -145.0 | |
| $S_{16}$ | $\text{ACA}_x$ | $2.21005 \cdot 10^{-4}$ | 34.3 | |
| $S_{17}$ | $\text{CN}_x^-$ | $5.52776 \cdot 10^{-3}$ | 163.1 | |
| $S_{18}$ | ATP | $1.90052 \cdot 10^{-3}$ | -2263.6 | |
| $S_{19}$ | ADP | $1.60722 \cdot 10^{-3}$ | -1388.7 | |
| $S_{20}$ | $\text{NAD}^+$ | $7.65889 \cdot 10^{-4}$ | -1128.9 | |
| $S_{21}$ | NADH | $2.14111 \cdot 10^{-4}$ | -1062.7 | |
| $S_{22}$ | AMP | $4.17601 \cdot 10^{-4}$ | -514.2 | |
| $S_{23}$ | product of reaction 22 | 0 | -2210.0 | $\mu^0$ and concentration set* |
| $S_{24}$ | $\text{Glc}_x0$ | $5.0 \cdot 10^{-3}$ | -392.3 | determines steady state |
| $S_{25}$ | $\text{CO}_2$ | 1 | -394.4 | concentration set** |
| $S_{26}$ | $\text{P}_i$ | 1 | -1055.5 | concentration set** |
| $S_{27}$ | $\text{CN}_x^-$ | $5.6 \cdot 10^{-5}$ | 163.1 | determines steady state |
| $S_{28}$ | $\text{H}_2\text{O}$ | 1 | -150.9 | concentration set** |
| $S_{29}$ | P, sink for $\text{ACA}_x$ | 0 | 34.3 | concentration set*** |
| $S_{30}$ | P, sink for $\text{EtOH}_x$ | 0 | 82.4 | concentration set*** |
| $S_{31}$ | P, sink for $\text{Glyc}_x$ | 0 | -145.0 | concentration set*** |
| $S_{32}$ | P, sink for $\text{Glyc}_x, \text{ACA}_x$ | 0 | 160.0 | concentration set*** |
| $S_{33}$ | substrate of reaction 24 | 1 | 0 | $\mu^0$ and concentration set**** |
| $S_{34}$ | product of reaction 24 | 1 | -895.0 | $\mu^0$ and concentration set**** |

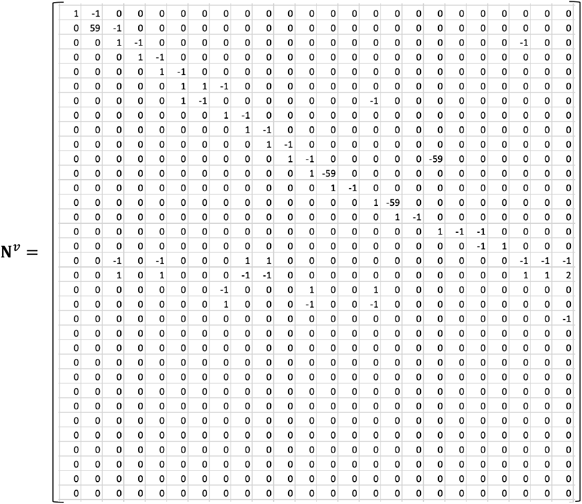

**Table 3:**
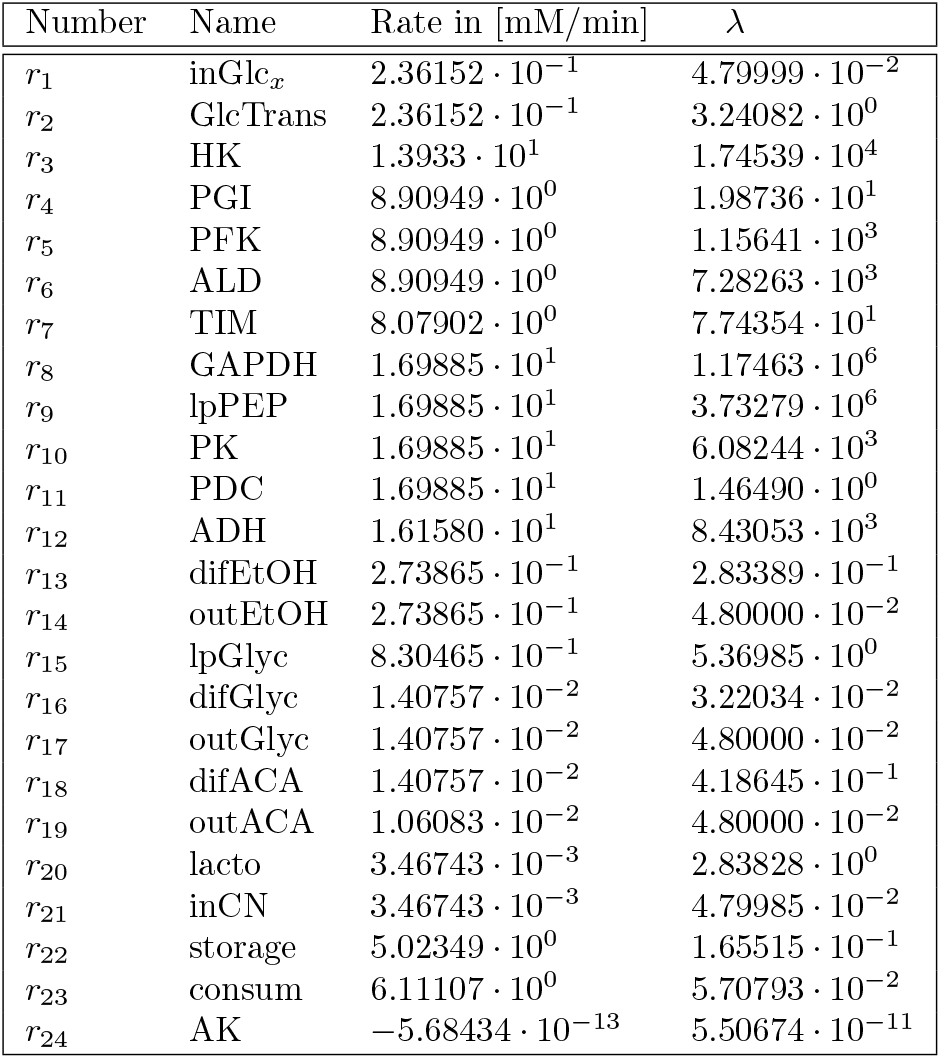
Reaction Information. Listed are all reactions of the model with their steady state rates and the respective *λ*-values. The unit of *λ*-values is dependent on the number of reactants and can vary between min^−1^ for one reactant, M^−1^ min^−1^ for two reactants, and M^−2^ min^−1^ for three reactants.

### 7.3 Derivation of *λ* -Values for a Linear Metabolic Pathway with Multiple Reactants per Reaction in Optimal States

The aim is to derive the values of *λ* in optimal states, i.e. states with given flux realized by minimal total enzyme concentration where the expression 30 holds. We consider again a linear pathway, but now each reaction can have one or several substrates and products as shown below

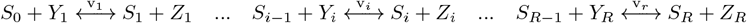

where *S*_0_ to *S*_*R*_ are the main or central series of metabolites of the pathway and 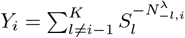 and 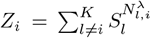 denote further substrates and products of reactions *v*_*i*_. Here and below, all equations hold for all *i* = 1, …*R*. Then, the rates read

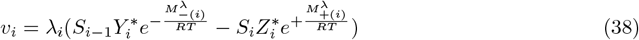

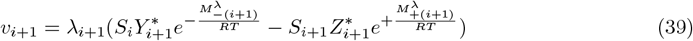

Using these rate expression, we will evaluate the expression

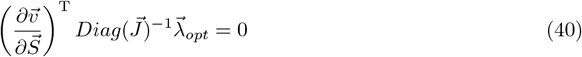

already introduced in Eq. 30. Here, the matrix 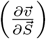 denotes the so-called elasticities, i.e. the partial derivatives of reaction rates w.r.t. metabolite concentrations, here to be evaluated at the optimal state. 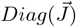 is the *R × R* matrix with steady fluxes in its main diagonal, resulting from the normalization used in Eq. 28. The expression 30 as such already establishes a relationship between *λ*-values and the kinetics of the network. Now, we need to evaluate how that relates to the dependency the logarithm of *λ*-values and the *M*-values. To this end, we use the TDE rate expressions, here exemplified for the *i*th and (*i* + 1)th reaction:

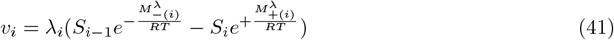

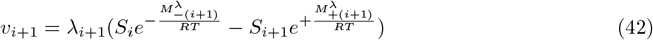

with 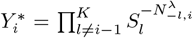 and 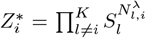. 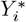 or 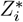 be 1 if the *i*-th reaction has only one substrate or one product, respectively. The elasticities then read

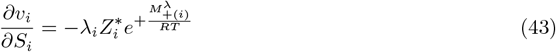

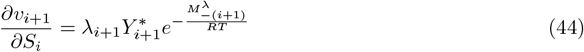

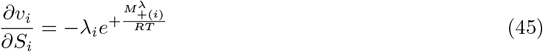

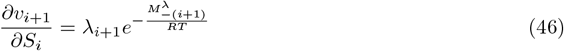

With these terms we can now do the matrix multiplication indicated in Equation 30. The *i*-th row (for the *i*-th substrate) reads as follows

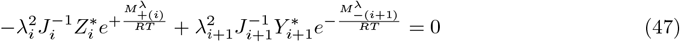

or

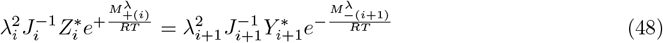

Note that the steady state flux through all reactions of a linear chain is the same, therefore, we can cancel *J*_*i*_ and *J*_*i*+1_. Hence, the ratio of two successive *λ*-values reads

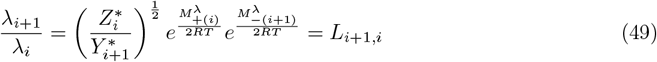

Together with the condition of a fixed (minimal) total amount of *λ*-values, i.e.

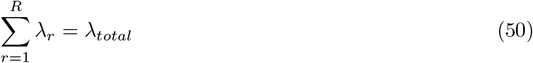

we easily find that

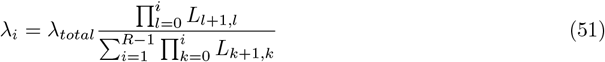

This expression establishes a systematic relationship between the *λ*-values for each reaction, an overall metabolic capacity of the system expressed by *λ*_*total*_ and the standard chemical potentials in the pathway.

### 7.4 Reaction Rates for Networks with only Monomolecular Reactions

In order to simplify the representation of reaction rates of a network in condensed form, we introduce the following abbreviation for the product of reactant concentration and the exponential of its standard chemical potential devided by *RT* as

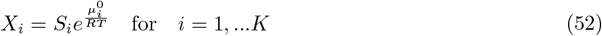

Using 52, the rate of a monomolecular reaction reversible converting *S*_*i*_ into *S*_*j*_ given in 7 reads

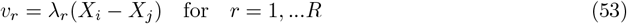

For a reaction network with the stoichiometric matrix **N** the vector of reaction rates, 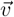 can be expressed as follows

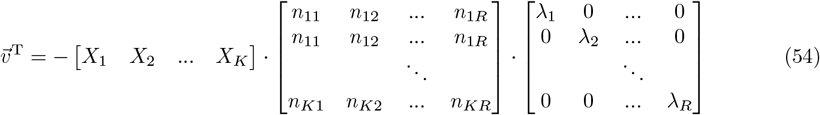

or

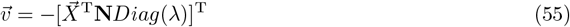

*Diag*(*λ*) denotes the *R × R* diagonal matrix containing the *λ* values in its main diagonal. In steady state, we have

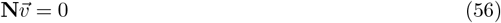

Introducing the vector for rates from (55) gives

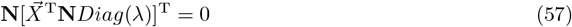

Rearranging yields

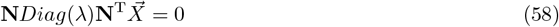

This is a homogeneous linear equation system, which could be solved either for the values of 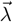 (i.e. the contribution of enzyme activities to rates) if 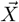 (i.e. the concentrations and standard chemical potentials of reactants) is given, or for the values of 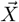 if 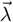 is given.

Let us now investigate the rates given by Equation 55 at the optimal state. First, we find that for the elasticities 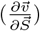 it holds

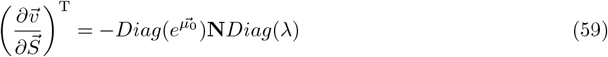

Multiplying with 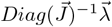 leads to

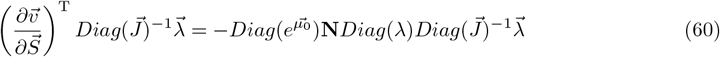

In the optimal state, using Equation 30, it follows

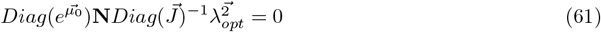

This is again a linear equation system for variables 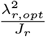 with *i* = *r*, …*R*. The matrix 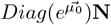 has the same size and rank as the stoichiometric matrix **N**, leading to the same number of independent solutions as for Equation 56. Using again the total amount of *λ* as given in Equation 50 then provides a scale for the values of the *λ*_*r*_.

### 7.5 Description of the *λ*-Value Optimization Algorithm

The algorithm is a multi-stage, iterative process designed to find the best possible “fit” for *λ*-values, particularly when dealing with incomplete data. It systematically evaluates input data, searches for optimal starting conditions, performs a guided optimization, and concludes with a comprehensive evaluation to select the most plausible parameter set. This process increases reliability and helps prevent convergence to suboptimal local minima. The graphical overview over the algorithm is provided in the flowchart 7.

The algorithm is broken down into four main phases:

#### 7.5.1 Phase 1: Initialization and Pre-calculations

This initial phase acts as a “triage” to determine the best starting point based on the provided dataset. The process follows one of two primary paths:

- **Case A: Sufficient Data** If the dataset is sufficient, a primary “fit” is calculated. The result is analyzed for quality, outliers, and gradient.
  - **Good Fit**: If the fit is good, the algorithm proceeds directly to the main optimization (Phase 3).
  - **Outlier Present**: If a small number of outliers are found, they are removed, and the fit is recalculated.
  - **Poor Fit**: If the fit is poor or untrustable, the algorithm proceeds to the fit-finding routine (Phase 2) to search for a better solution.
- **Case B: Insufficient Data** If the data is too sparse for an initial fit, a “naive” optimization is performed to get a basic starting point.
  - If this result “seems to be promising,” i.e. all criteria are fulfilled, it is presented to the user for manual validation.
  - If not, the algorithm proceeds to the fit-finding routine (Phase 2).

**Figure 7.**
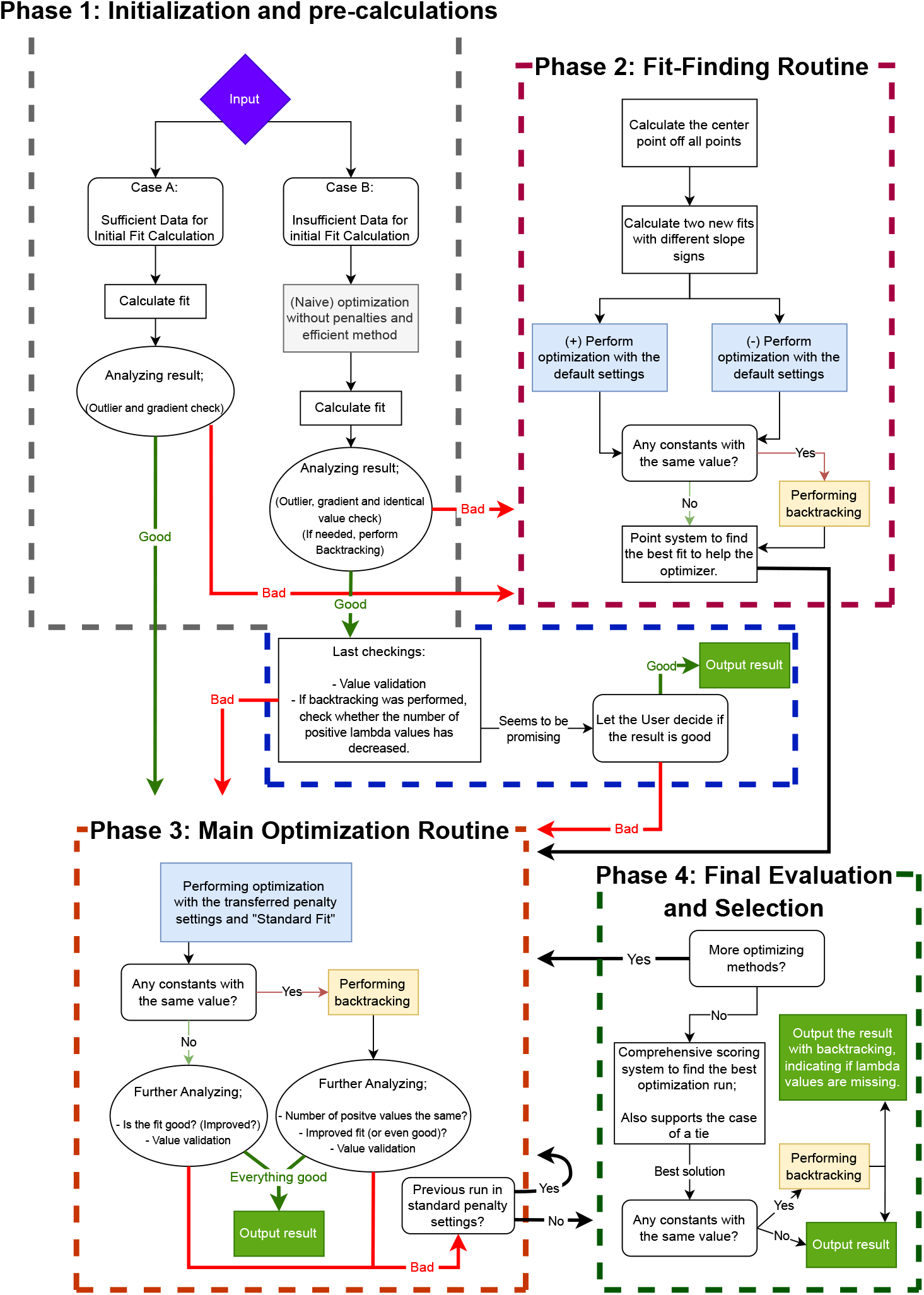
Overview of the Optimization Algorithm.

#### 7.5.2 Phase 2: Fit-Finding Routine

This phase is a systematic “fit-seeker” that activates when the initial data quality is insufficient. Its goal is to establish a robust foundation for the main optimization.

1. **Center Point Calculation**: It begins by calculating the center point of the most trustworthy data points available.
2. **Hypothesis Generation**: Based on this center, two competing “fit” hypotheses are generated (e.g., one with a positive slope and one with a negative slope).
3. **Optimization & Selection**: A brief, independent optimization is run for both hypotheses. A quantitative scoring system compares the two results, and the most promising candidate is selected to be passed to the main optimization phase.

#### 7.5.3 Phase 3: Main Optimization Routine

This is the core computational engine of the algorithm, designed to refine the starting fit into a final, optimized solution.

1. **Standard Optimization**: The process begins with an optimization run using standard, default parameters.
2. **Anomaly Detection**: The resulting solution is analyzed for potential issues, such as:
  - **Calculation Errors**: Checking for invalid values, like ‘NaN’ (Not a Number) concentrations.
  - **Degeneracy**: Checking if many *λ*-values have converged to the same identical value.
3. **Backtracking**: If such anomalies are detected (e.g., identical *λ*-values), a “backtracking” procedure is initiated to compute the true *λ*-values from the found concentrations and refine the solution.
4. **Recursive Refinement**: If the result remains suboptimal after this analysis, the routine calls itself recursively, this time applying **higher penalty constraints** to compel the optimizer toward a more robust solution.

#### 7.5.4 Phase 4: Final Evaluation and Selection

This final phase acts as the “ultimate judge”, evaluating all solutions generated during the entire process to select the single best result.

1. **Comprehensive Scoring**: A point system scores every fit. Points are awarded or penalized based on several criteria, including:
  - The **coefficient of determination** (*R*^2^).
  - The **distribution of** *λ***-values** (penalizing identical values).
  - The **slope** of the fit.
2. **Tie-Breaking**: A multi-stage tie-breaking protocol is implemented to assign penalty points for undesirable traits and ensure a unique winner is chosen.
3. **Final Verification**: The selected solution may undergo a final backtracking procedure before being output as the definitive solution.

### 7.6 List of Example Models Analyzed for this Study

Below we present a visual summary of the results for further analyzed published models. All values for concentrations, standard chemical potentials, fluxes and resulting *λ*-values can be found in the Supplementary Information “Results Model Analysis.xlsx”.

**Figure 8.**
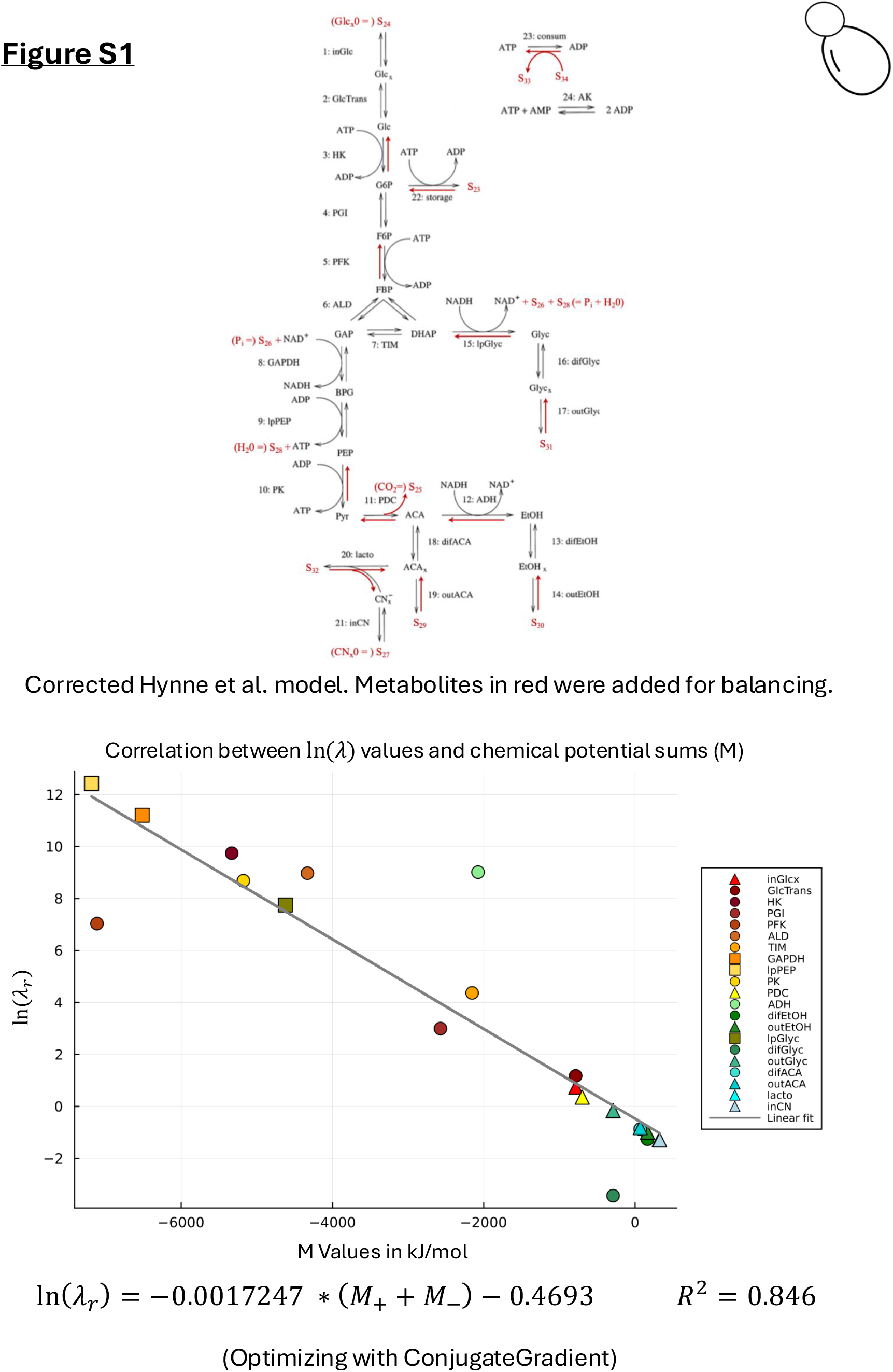
Optimized Hynne Model

**Figure 9.**
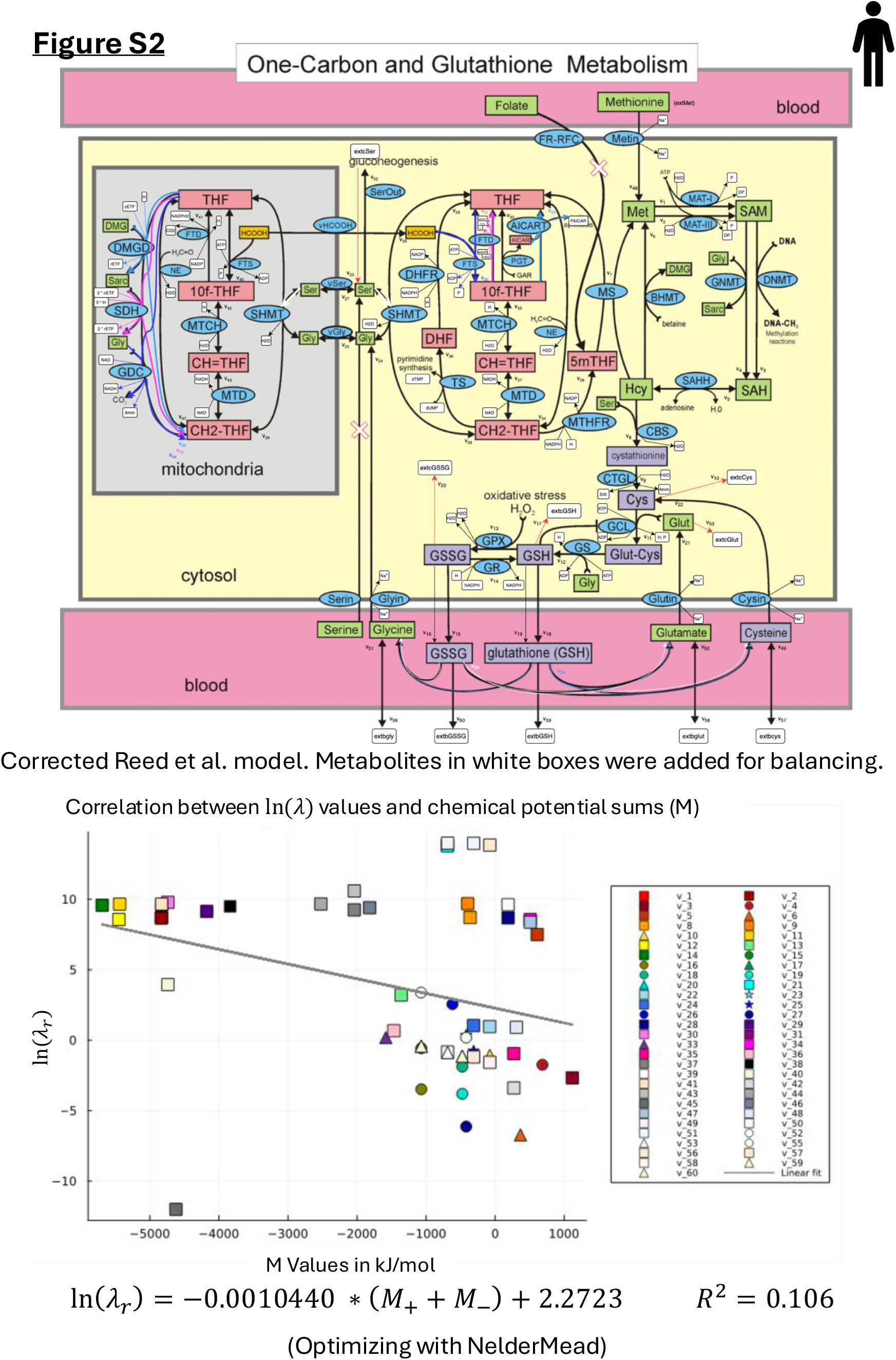
Optimized Reed Model

**Figure 10.**
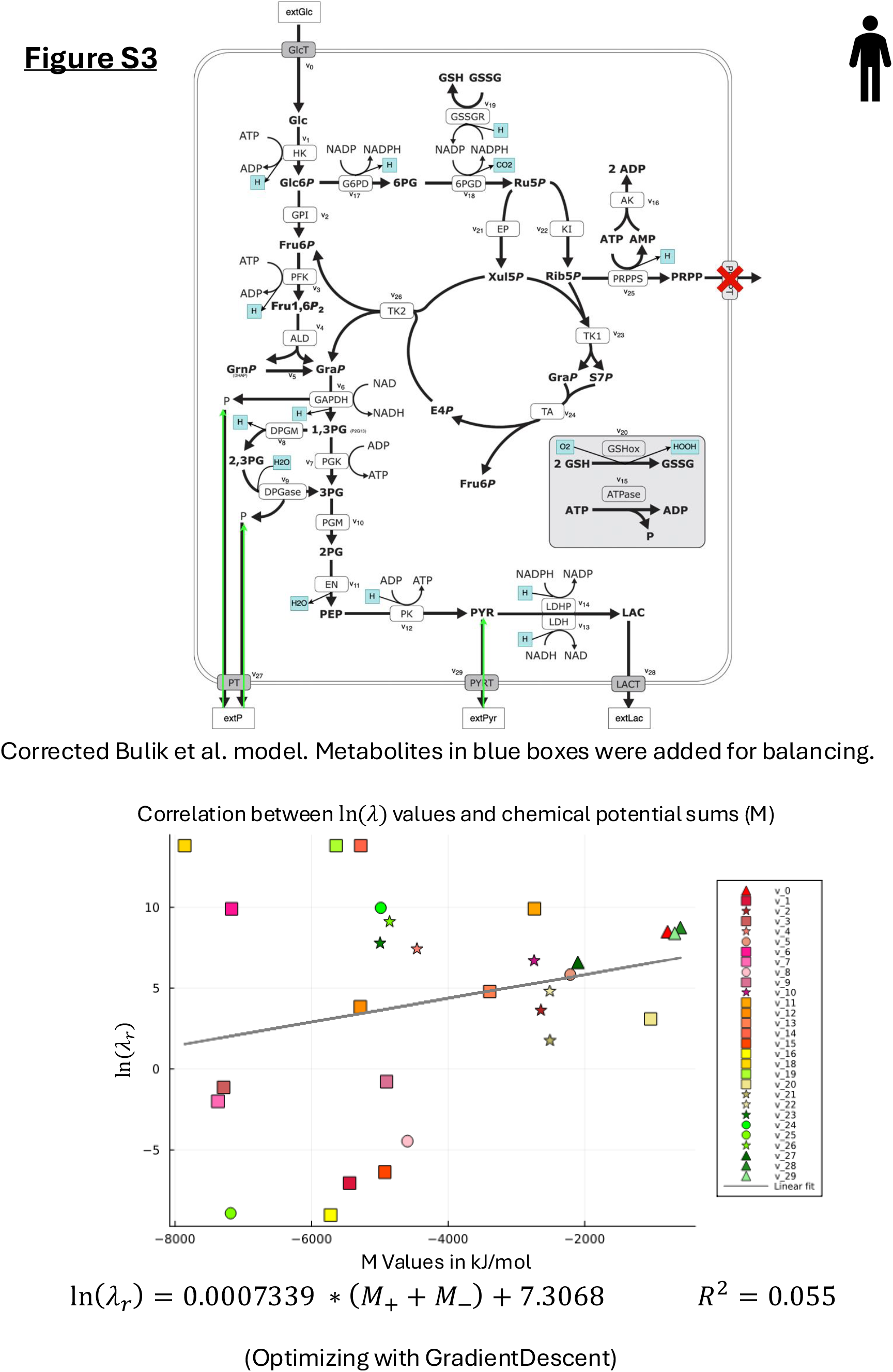
Optimized Bulik Model

**Figure 11.**
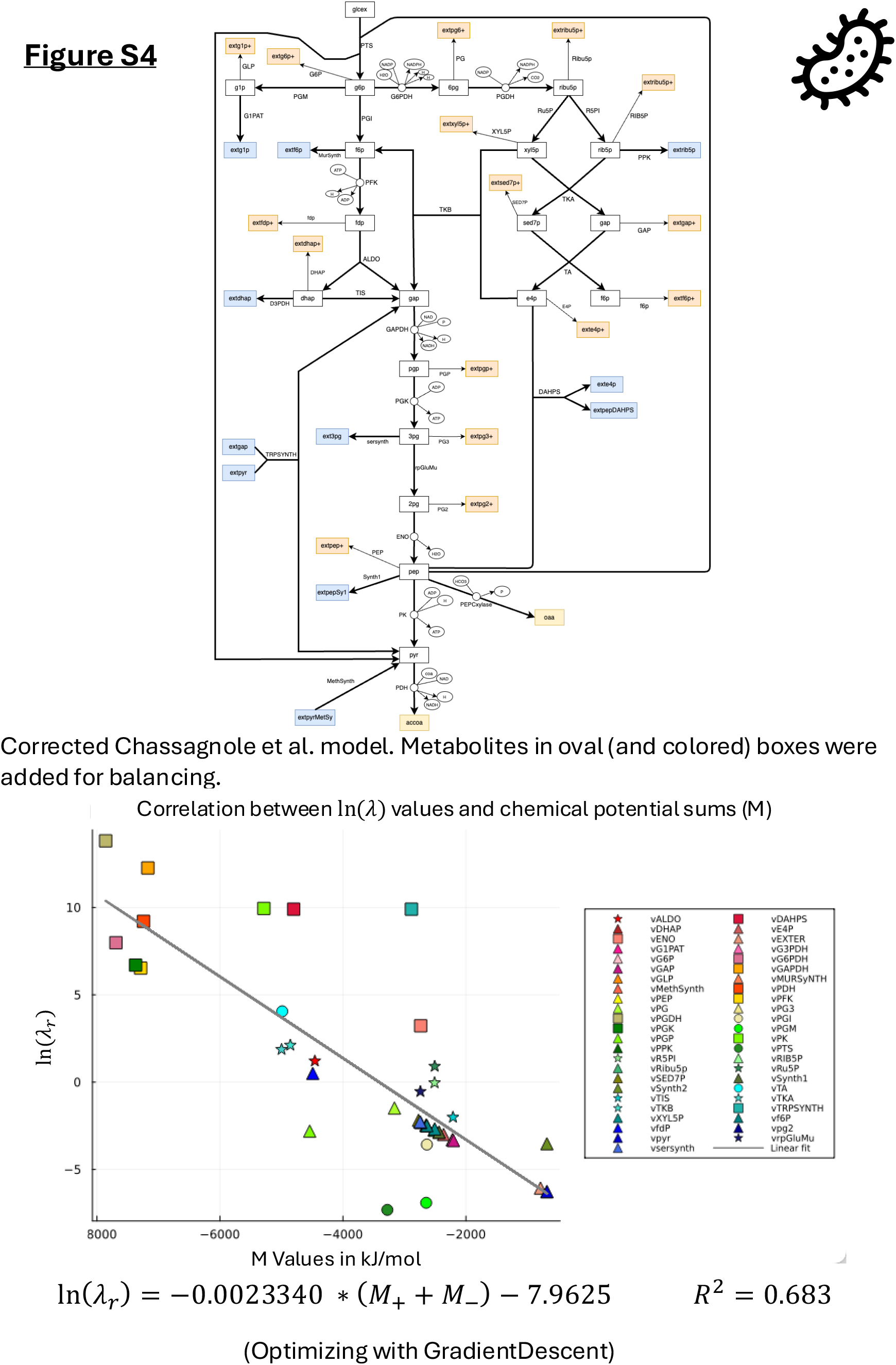
Optimized Chassagnole Model

**Figure 12.**
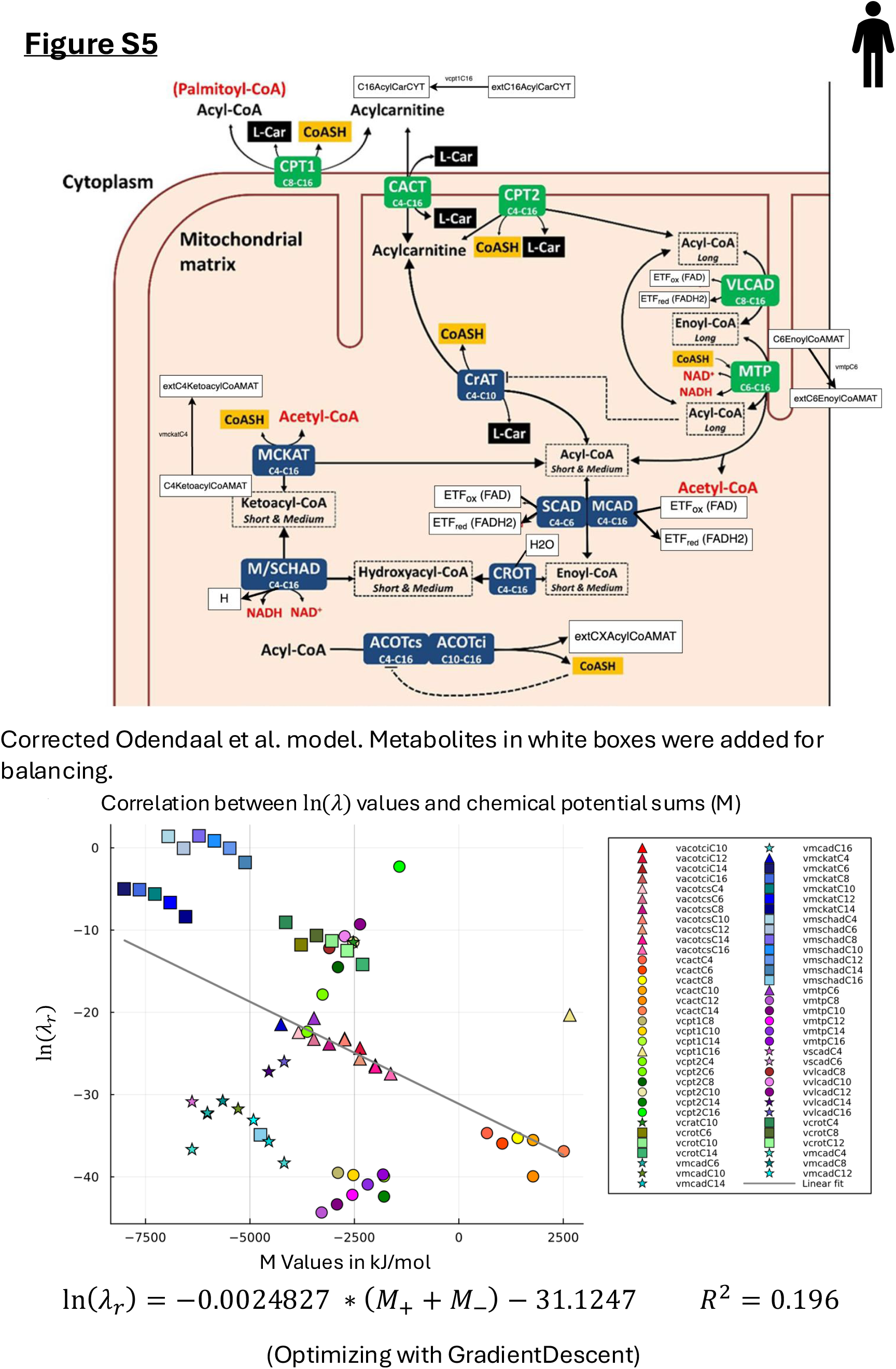
Optimized Odendaal Model

**Figure 13.**
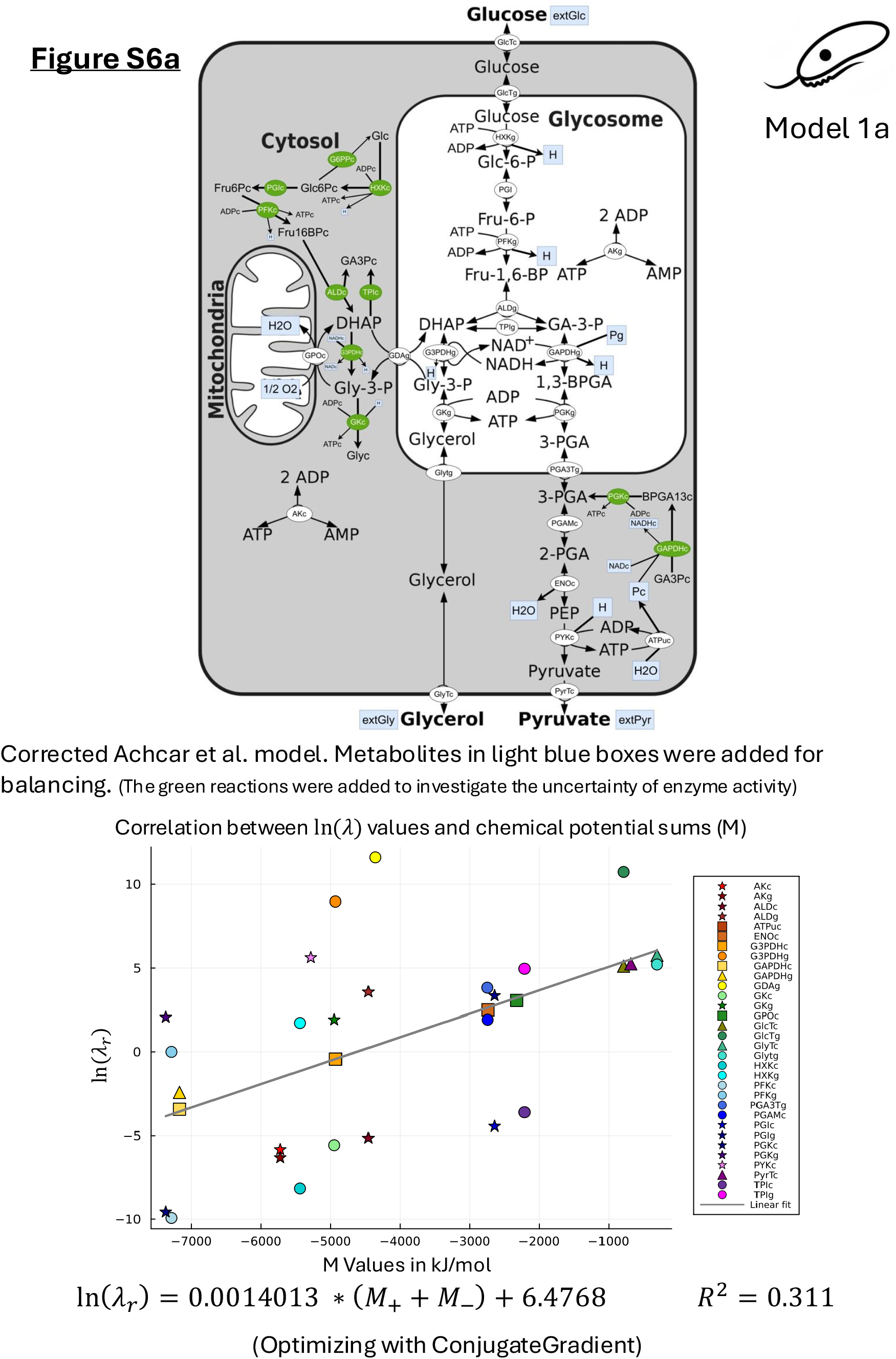
Optimized Achcar Model, Version 1

**Figure 14.**
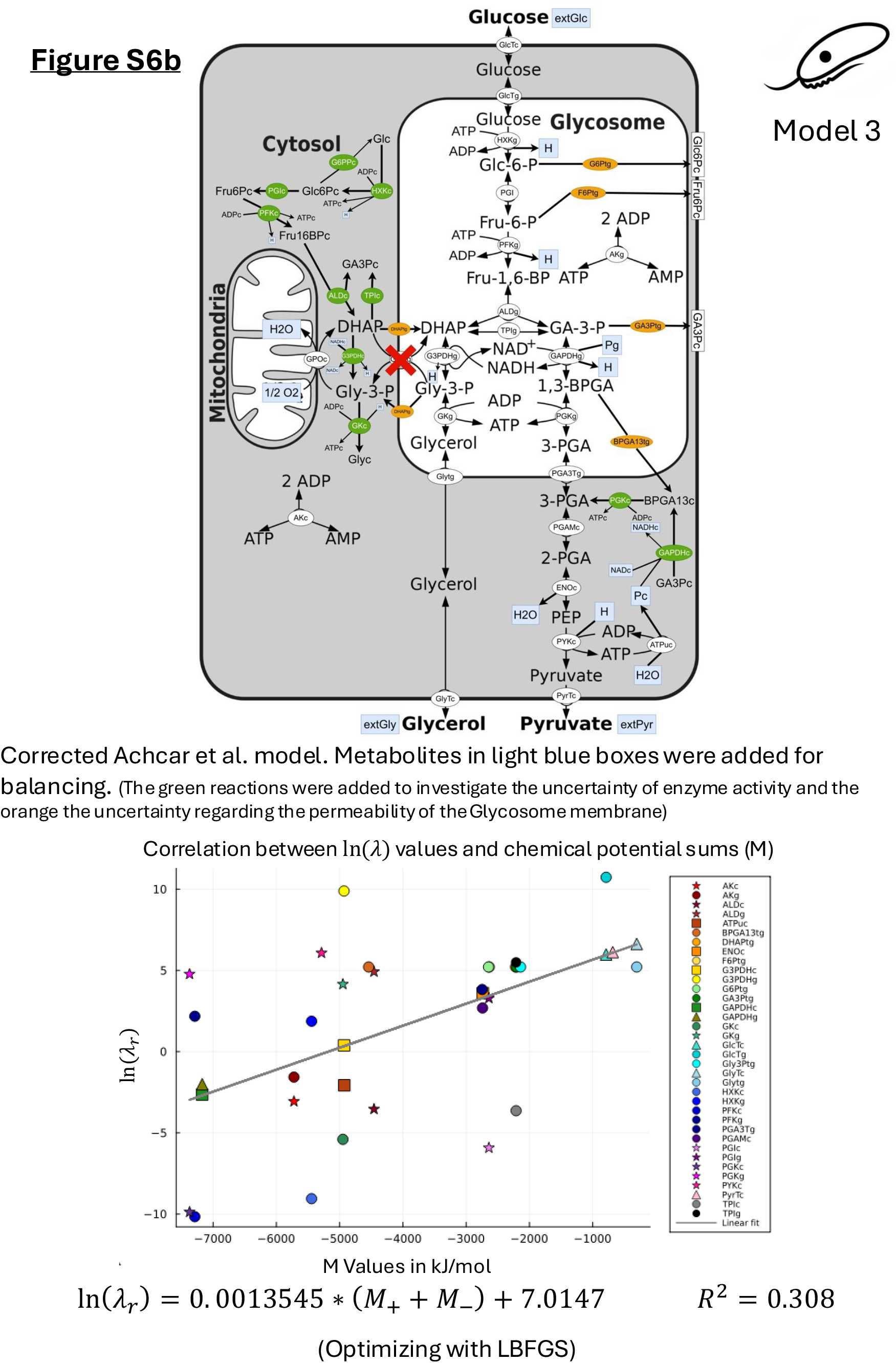
Optimized Achcar Model, Version 2

**Figure 15.**
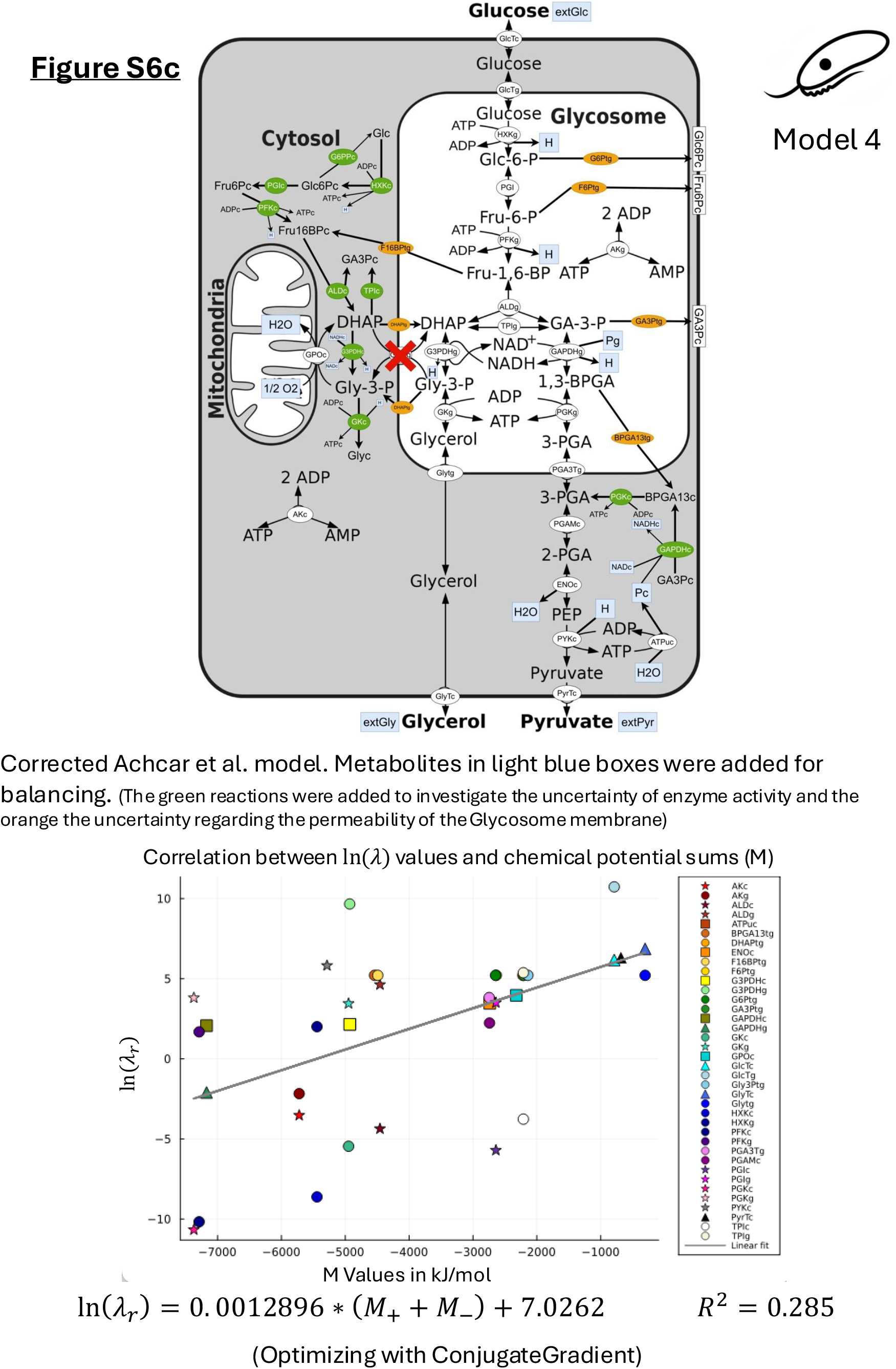
Optimized Achcar Model, Version 3

## Notes

### Competing Interest Statement

The authors have declared no competing interest.

### Summary of Updates

A justification for the found correlation between standard chemical potentials and kinetic capacity factor has been added. An optimization algorithm to determine the kinetic capacity factors for networks with partially known concentrations and fluxes was added.

